# Endoglin deficiency elicits hypoxia-driven congestive heart failure in zebrafish

**DOI:** 10.1101/2021.10.09.463775

**Authors:** Etienne Lelièvre, Charlotte Bureau, Yann Bordat, Maxence Frétaud, Christelle Langevin, Chris Jopling, Karima Kissa

## Abstract

Hereditary hemorrhagic Telangiectasia (HHT) is a rare genetic disease relying on mutations affecting components of Bone Morphogenetic Protein and Transforming Growth Factor-β (BMP/TGF-β) signaling pathway in endothelial cells. This disorder is characterized by arterio-venous malformations prone to rupture. and ensuing hemorrhages are responsible for iron deficiency anemia. Along with Activin receptor-like kinase ALK1, Endoglin is involved in the vast majority of HHT cases. In this report, we characterized zebrafish endoglin locus and demonstrated that it produces two phylogenetically conserved protein isoforms using a distinctive alternative splicing mechanism. Functional analysis of a Crispr/Cas9 zebrafish Endoglin mutant revealed that Endoglin deficiency results in massive death during the course from juvenile stage to adulthood. Endoglin deficient fish develop a cardiomegaly resulting in heart failure and hypochromic anemia which both stem from chronic hypoxia. Histological analysis and confocal imaging evidenced structural alterations of the developing gill and its underlying vascular network that tally with hypoxia. Finally, phenylhydrazine treatment demonstrated that lowering hematocrit/blood viscosity alleviates heart failure and enhances survival of Endoglin deficient fish. Altogether, our data indicate that Endoglin is crucial for gill vascular development and that further studies using zebrafish in general and this endoglin mutant in particular will provide crucial hints regarding the molecular and cellular events altered in HHT for the development of new therapeutic strategies.

**Summary Statement:** Endoglin deficiency in zebrafish recapitulates critical aspects of Hereditary Hemorrhagic Telangiectasia (HHT) and will thus constitute a valuable model in large scale screens for HHT-active drugs.

## Introduction

Endoglin (CD105) belongs to the Bone Morphogenetic Protein and Transforming Growth Factor-β (BMP/TGF-β) receptor superfamily and binds TGF-β1, TGF-β3, BMP9 and BMP10 (Castonguay et al., 2011; Cheifetz et al., 1992). Like betaglycan (TGBR3), this single-pass transmembrane protein works as an ancillary receptor by modulating the signaling activity of the heterotetrameric receptor complex constituted of type I and type II serine-threonine kinase receptors which phosphorylate R-Smads which eventually associates with Co-Smad (Smad4) to form a complex that regulates the transcription of target genes via Smad Binding Elements (SBE). Endoglin is mainly expressed in endothelial cells (EC) and Endoglin deficient mice die around E10.5 from cardiovascular defects associated with improper vascular smooth cell coverage (Li et al., 1999). In addition to endothelial cell expression, in mammals, Endoglin is also expressed in neural crest cells and derivatives such as vascular smooth muscle cells responsible for blood vessel stabilization and vascular tone control and in Hematopoietic Stem cells (HSCs) as they emerge from intraaortic clusters to later being restricted to HSCs with long term repopulating capacities (LT-HSCs) (Chen et al., 2002; Mancini et al., 2007).

In humans, inactivating mutations in endoglin (ENG) gene are responsible for type 1 Hereditary Hemorrhagic Telangiectasia (HHT) while HHT2 is due to mutations affecting ACVRL1 (ALK1) a type I TGF-β receptor specifically expressed in endothelial cells. Together, HHT1 and HHT2 account for approximately 85% of diagnosed HHT cases (McDonald et al., 2015), for review. Endoglin and ALK1 independent HHT cases rely on mutations affecting protein function of BMP9, a high affinity ligand for ALK1, microprocessor RNAse III Drosha and additional uncharacterized loci while mutations in SMAD4 are causative of Juvenile Polyposis-HHT (Gallione et al., 2006; Jiang et al., 2018; Wooderchak-Donahue et al., 2013).

Also known as Rendu-Osler-Weber syndrome, HHT is a rare inherited autosomal dominant genetic disorder characterized by a wide array of symptoms among which mucocutaneous telangiectasias and recurrent epistaxis are the foremost diagnosis cues. These angiodysplastic lesions or arteriovenous malformations (AVMs) also often affect organs such as the gastrointestinal tract, the lungs, the liver and the brain. They consist of dilated veins prone to rupture under mechanical strains. These AVMs resulting from the loss of the arteriolar-capillary plexus connect venous vessels directly to arteries exposing the former to arterial-type blood flow mechanics (for review (Guttmacher et al., 1995)).

Although much concern arise from brain, liver or lung AVMs which can have life-threatening outcomes, massive blood loss from hemorrhages affecting the gastrointestinal tract or from recurrent epistaxis are also important issues as it results in iron deficiency anemia that requires medical management by way of regular iron replenishing cures and/or blood transfusion (Stross, 2013).

Congestive heart failure constitutes another critical clinical manifestation of HHT although considered fairly rare. Cases seem to be mostly associated with liver AVMs and chronic anemia of iron deficiency type (Cho et al., 2012; Goussous et al., 2009; Montejo Baranda et al., 1984; Wu et al., 2017)

Albeit different in terms of incidence in specific organs such as brain or liver which is much higher in HHT1, all HHT forms rely on mutations affecting genes working on a common TGF-β/BMP pathway and recent findings regarding Drosha indicate that non-classical TGF-β/BMP pathways are also involved (Letteboer et al., 2006). Thus, while this pathway is indisputably at the core of the pathology, tissue specific cues or vascular bed specificities might provide a terrain that will somehow trigger or favored the onset of the pathology. In humans, HHT relies on hemizygous mutations and accordingly, mice carrying a single knockout Endoglin allele do spontaneously develop HHT-like phenotypes with varying penetrance depending on genetic background (Bourdeau et al., 1999). This collectively refers to as the “second hit” concept by which additional alterations would be required for the development of the pathology. Consistent with this concept, angiogenic/inflammatory environments have been found to be potent drivers of AVM formation in both heterozygous knockout Endoglin and ALK1 mouse models (Tual-Chalot et al., 2015).

Several recent studies using BMP9/10 blocking antibodies, inducible tissue specific Endoglin, ALK1 and Smad4 knockout mouse models as well as Endoglin genome editing in zebrafish have provided important insights into the cellular and molecular mechanisms governing AVM formation which involves inappropriate signaling responses to blood flow including EC polarization, venous/arterial identity maintenance, proliferation and pericyte recruitment (Baeyens et al., 2016; Jin et al., 2017; Ola et al., 2016; Ola et al., 2018; Sugden et al., 2017). In keeping with this, in HUVECs, laminar shear stress triggers Endoglin association with ALK1 and potentiates ALK1 signaling pathway in response to endogenous serum BMP9/10 concentration (Baeyens et al., 2016).

In this work, we examined the consequences of Endoglin loss in postembryonic stage zebrafish. Using a CRISPR/Cas9 endoglin mutant, we demonstrated that homozygous mutants massively die from congestive cardiomyopathy from about 1 month old accompanied with iron deficiency anemia. We found that this pathological condition sets just before 15dpf when hypoxia and cardiac stress markers start increasing in mutants and coincide with heart chamber enlargement which involves cardiomyocyte proliferation. Data from histology and imaging of blood vessel organization during gill development strongly suggest that hypoxia stems from improper gill functioning due to Endoglin direct activity in this organ. Controlling hematocrit/blood viscosity triggered by chronic hypoxia markedly lessens hypoxic response, cardiac stress to result in enhanced survival of Endoglin deficient fish. Thus, by reproducing important features of HHT, our zebrafish model of Endoglin deficiency will provide the ground for future detailed molecular analysis that will be important for both the identification of HHT altered signaling pathways and the development of new treatments.

## Results

### Zebrafish endoglin locus, transcripts and expression characterization

To develop tools to address Endoglin expression and function in zebrafish, we first characterized *endoglin* locus and associated transcripts in this organism. We conducted 5’ and 3’ RACE experiments and obtained complete and trustful sequences. This allowed to reconstruct exon-intron organization of zebrafish endoglin gene, essential for gross mapping of transcriptional regulatory regions. We identified 3 previously undescribed exons (Fig. 1A). Two were non-coding exons, one very short located the utmost 5’ positioning Endoglin Transcription Start Site and one 3’ containing most of Endoglin 3’UTR. The third identified exon (exon 13 in our nomenclature) corresponded to an alternate exon which introduces an early termination codon resulting in an Endoglin protein isoform with a very short cytosolic domain (Fig. 1B and Fig. 1C), an isoform also expressed in mammals (Bellón et al., 1993). By contrast to Long isoforms, phylogenetic alignments revealed little or no residue conservation in the cytosolic tail of Short isoforms (Fig. 1D). RT-PCR analysis demonstrated that both variants were expressed in embryonic and adult tissues, endoglin messenger for the short isoform being markedly less abundant (Fig. 1B). Similar analysis performed on developmentally staged embryos indicated that with exception of a marginal maternal contribution, the zygotic expression of endoglin coincided with early somitogenesis and gradually expanded in later stages with a somewhat stable Long isoform/Short isoform ratio (Fig. 1E). Whole-mount *in situ* hybridization experiments detected endoglin expression in developing blood vessels with the highest endoglin expression mostly associated with veins (Fig. 1F). In line with early findings in mammals, our data also confirmed the results from a previous study in zebrafish (Sugden et al., 2017) and suggested of a conserved Endoglin function throughout the whole vertebrate phylum.

**Fig. 1.**
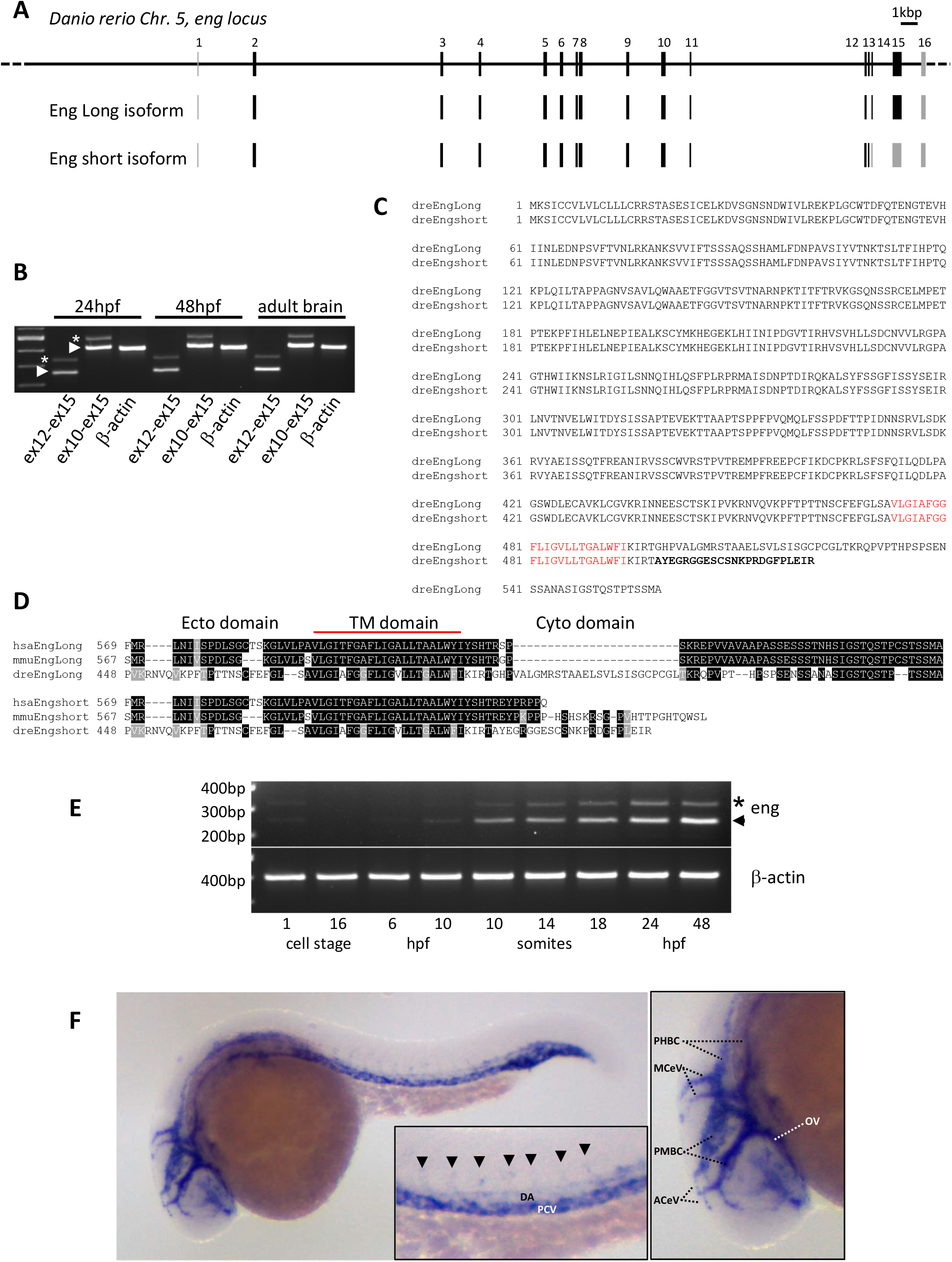
Characterization of zebrafish Endoglin chromosomal organization, transcripts and expression pattern. (A) Zebrafish *endoglin* gene exon-intron structure on chromosome 5. Coding exons (black), non-coding exons (grey). (B) RT-PCR analysis of endoglin variants. Endoglin Long (arrowhead) and short (asterisk) isoform messengers expression in embryo and adult tissue (brain). (C) Protein alignment of zebrafish Endoglin isoforms. Alternate amino acids (aa) of Endoglin short isoform are in bold. Transmembrane aa (red). (D) Alignment of Human, mouse and zebrafish Endoglin short isoform. Identical (black) and similar (grey) aa. (E) RT-PCR analysis of endoglin variants during zebrafish early development. Endoglin short (asterisk), Endoglin Long (arrowhead) isoform. (F) Whole-mount in situ hybridization using dig-labelled endoglin antisense riboprobe on 24hpf wild-type embryo. Inset, close-up of trunk region showing endoglin differential expression between dorsal aorta (DA) and posterior cardinal vein (PCV). Intersegmental vessels (arrowheads). Right panel, Close-up of head region showing mostly venous endoglin expression, ACeV, Anterior cerebral vein; MCeV, Middle cerebral vein; PHBC, Primordial hindbrain channel; PMBC, Primordial midbrain channel; OV, optic vein

### CRISPR/Cas9 generation of endoglin mutant zebrafish

To address Endoglin function in zebrafish, we used a CRISPR/Cas9 approach based on a guide RNA overlapping with endoglin ATG located on exon 2. To avoid potential deleterious effects of Endoglin inactivation such as observed in mouse, fish from 2 different injection setups (1-2 cell-stage (early) versus 4-16 cell-stage (late)), intended that late-stage injection would result in higher mosaicism and survival, were raised to adulthood. Analysis of germline presence of indels in sperm samples or in clutches from mating with wild type fish detected indels in 18% (2/11) and 94% (32/34) of fish resulting respectively from early and late injections indicating that late-stage injection indeed circumvented lethality. Interestingly, numerous fish did not recover from anesthesia/sperm sampling and massive bleeding from the gills was systematically observed in these fish. By contrast with fish recovering normally from the sampling procedure, bleeders were indel carriers. We, however, managed to obtain a single clutch from one male which died during re-sampling from which we derived the line analyzed in this study. This founder (F_0_) transmitted to 26.4% of the progeny (27/102) a unique indel consisting in a 2bp deletion and a C>A or T>A base change thus destroying endoglin translation initiation ATG codon and nucleotide −1 and −2 of endoglin Kozak sequence (Fig. 2A). Among F_1_, some fish later found to carry a mutated allele died at a juvenile stage (3-4 weeks) and some other showed overt redness later on but the vast majority developed normally and were indiscernible from wild-type siblings. Taken together these data suggested that endoglin mutation would be correlated with late development and/or survival issues in zebrafish.

**Fig. 2.**
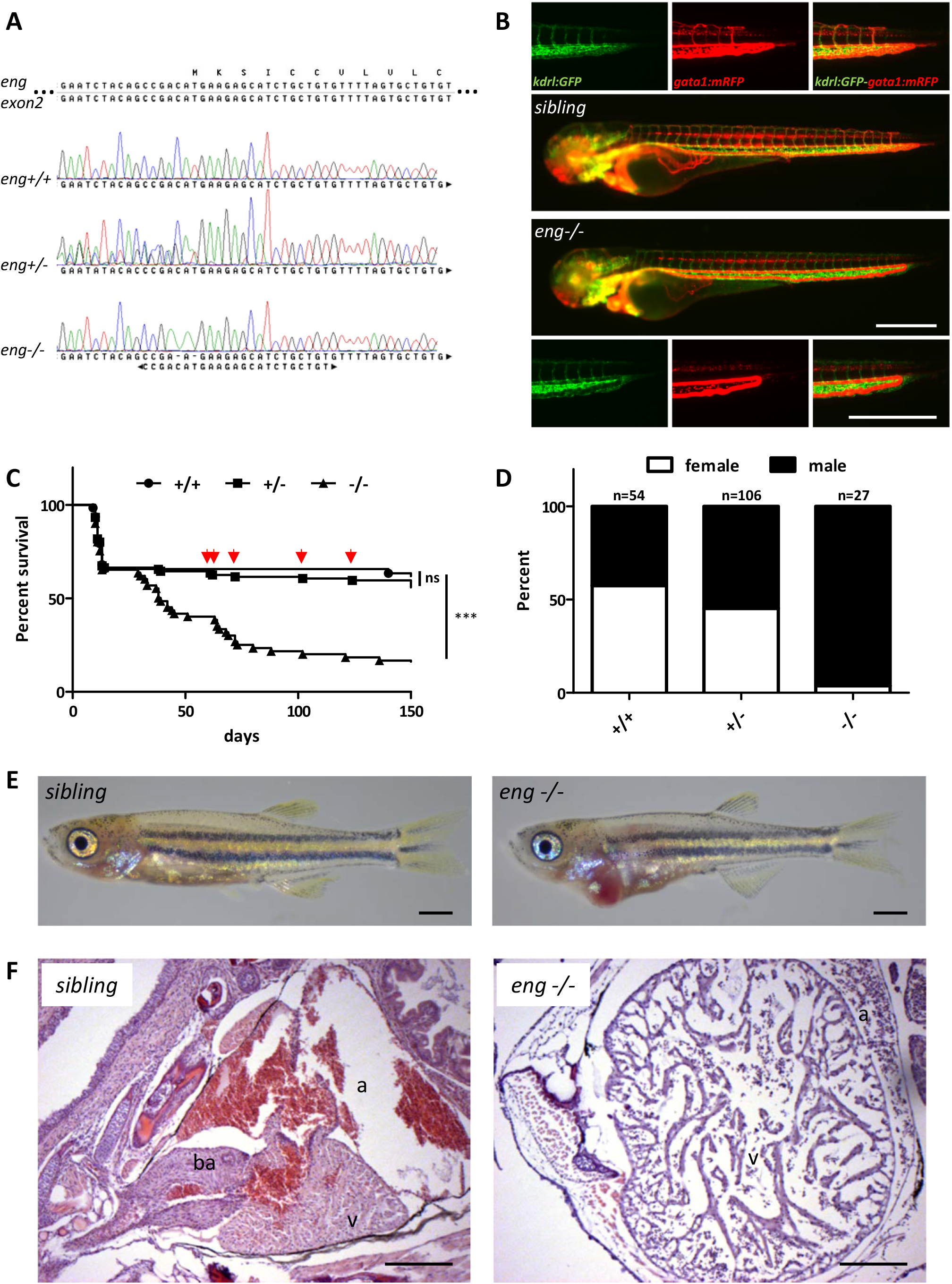
Endoglin deficiency results in congestive heart failure in zebrafish. (A) Sanger sequencing chromatograms of gRNA targeted region on endoglin exon 2 in wild-type (*eng*^*+/+*^), heterozygous (*eng*^*+/−*^) and homozygous (*eng*^*−/−*^) mutants. Complementary gRNA sequence is indicated below chromatograms. (B) Imaging of 72hpf *eng*^*−/−*^ and siblings blood flow pattern. Analysis performed in *Tg(kdrl:GFP)*, *Tg(gata1:mRFP)* background to highlight blood vessels and erythrocytes respectively. Pictures shown are representative of data from siblings (n=10) and *eng*^*−/−*^ (n=8). Bar, 500 μm. (C) Kaplan-Meyer representation of *eng*^*+/+*^, *eng*^*+/−*^ and *eng*^*−/−*^ survival. +/+ vs −/− and +/− vs −/− *P*<0.0001, +/+ vs +/− ns, not significant in Log-rank (Mantel-Cox) Test. +/+ n=45, +/− n=104, −/− n=56. Arrows point to symptomatic siblings later identified as *eng*^*+/−*^. (D) Influence of genotype over sex ratio in 3 months plus individuals. Data from two complete independent experiments plus two focused on *eng*^*−/−*^. Total number of individuals analyzed is indicated above bars. (E) Representative morphology of 30dpf *eng*^*−/−*^ and siblings. Note the enlarged cardiac area and overall paleness in *eng*^*−/−*^ fish. Bar, 1 mm. (F) Hematoxyllin-eosin staining of heart histological sections reveals dramatic enlargement and structural alteration of the ventricule, hypochromic red blood cells and swollen surrounding tissue. Bar, 100 μm. (a, atrium; ba, bulbus arteriosus; v, ventricule)

### Endoglin deficient zebrafish massively die from congestive heart failure

Above observations along with Endoglin established role in HHT and midgestation lethality in homozygous knockout mice prompted us to analyze the effect of endoglin mutation in incrosses of heterozygotes fish. By 3dpf, we observed a strong modification of blood flow pattern in a dilated dorsal aorta (DA) – Posterior cardinal vein (PCV) loop and a poor perfusion of Intersegmental vessels (ISVs) (Fig. 2B). RT-qPCR demonstrated that endoglin messenger abundance was reduced by 36.6% and 26.2% in homozygous mutants (*eng*^*−/−*^) compared to wild-type and siblings, respectively, which could reflect nonsense mRNA decay induced by deleterious mutation (Supplementary Fig. S1A). Altogether our results confirmed earlier report (Sugden et al., 2017) and strongly suggested that the mutation we created also corresponded to a null allele. Endoglin mutation effect was evaluated in survival experiments from 3dpf onwards. Kaplan-Meier survival plots showed a high mortality rate of *eng*^*−/−*^ fish (Figure 2C). Indeed, while 63.5% of *eng*^*+/+*^ and 59.6% of *eng*^*+/−*^ survived up to 5 months only 16.8% of *eng*^*−/−*^ managed to reach this age. Data indicated that *eng*^*−/−*^ started to die massively by the age of 1 month-old and displayed a median survival of 44 days. A fraction of *eng*^*−/−*^ fish survived the 5 months period but survival figures underestimated the severity of the phenotype as over half (7/13) of *eng*^*−/−*^ surviving fish never reached adulthood and stalled in a 11-17mm length range that precluded macroscopic gender determination. Surprisingly, *eng*^*−/−*^ individuals reaching adulthood were found to be almost exclusively phenotypic males (Fig. 2D). Because zebrafish sex is not assigned chromosomally, this could reflect either a genuine bias towards males or a higher sensitivity of females to Endoglin deficiency. Examination of *eng*^*−/−*^ at 30dpf i.e. the very onset of lethality revealed that most fish exhibited a red enlarged cardiac area whereas the rest of the body appeared paler than siblings (Fig. 2E). We also noticed that, by contrast with siblings, *eng*^*−/−*^ were hyperventilating and displayed a marked surface respiratory behavior (SRB). To get a better insight into the cardiac issue, we analyzed histological sections of the cardiac region (Fig. 2f). From these, the ventricule of *eng*^*−/−*^ appeared dramatically oversized. Red blood cells in *eng*^*−/−*^ did not present the characteristic orange color resulting from eosin reaction with hemoglobin, seen in siblings thus revealing a decrease in hemoglobin content. Surrounding connective tissue also appeared loose indicating that edema, an accompanying symptom of heart dysfunction, was also taking place. Altogether this data demonstrated that Endoglin deficient fish developed congestive heart failure as well as anemia with lethal outcome.

### A single endoglin mutant allele is sufficient to induce pathology

Based on macroscopic analysis, 3 month old or older *eng*^*−/−*^ surviving fish fell into 3 classes of phenotype severity: 1) fish with arrested growth, hyperventilating and with recurrent enlarged cardiac area, 2) adult size fish hyperventilating with enlarged cardiac area to various degrees, 3) fish macroscopically asymptomatic (Supplementary Fig. S2). Interestingly, a fraction of *eng*^*+/−*^ was also found to be symptomatic i.e. hyperventilating with occasionally enlarged cardiac area (Supplementary Fig. S2). Symptomatic *eng*^*+/−*^ were observed around one month old and raised apart to monitor the evolution of phenotypes and life expectancy. Collective data from two independent experiments indicated that the frequency of symptomatic *eng*^*+/−*^ averaged 14% ± 3% of total *eng*^*+/−*^ (16/114). In addition to hyperventilation and SRB, their most striking feature was their marked ruddy complexion which persisted as they completed growth to adulthood as well as the occasional presence of dilated surface vessels reminiscent of telangiectasias (Supplementary Fig. S3). Although the number of such animals was too low to significantly affect heterozygotes overall survival rate, we did observe a higher frequency of death events in this group (Fig. 2C, arrowheads) resulting in an estimated 5-month survival of 37.5% (6/16). Thus, similar to HHT1 mouse model, a single mutated endoglin allele resulted in pathological conditions with limited penetrance and were in line with the “second hit” concept, in zebrafish as well.

### Early detection of concomitant hypoxic and cardiac stress responses in Endoglin deficient zebrafish

Hypoxia being a potent driver of cardiac remodeling involving cardiomyocyte hypertrophy in mammals and both proliferation and hypertrophy in zebrafish (Jopling et al., 2012; Ke et al., 2017; Sun et al., 2009), we reasoned that it might trigger heart failure in our model. Supporting this hypothesis, hyperventilation and SRB appeared as characteristic manifestations in *eng*^*−/−*^. First, we monitored the expression of cardiac stress markers, i.e. atrial- and brain-type natriuretic peptides *nppa* and *nppb* respectively, to define when the heart of *eng*^*−/−*^ started being at stake. Using RT-qPCR, we found that by 15dpf, when compared to siblings, *eng*^*−/−*^ exhibited a 1.8 and a 3.3 fold induction of *nppa* and *nppb* respectively. These differences in expression between *eng*^*−/−*^ and siblings then expanded to reach up to 49 and 223 fold change of *nppa* and *nppb* abundance respectively, consistent with the gradual enlargement of *eng*^*−/−*^ heart region over time (Fig. 3A). We, thus, analyzed the expression of well accepted hypoxia-responsive genes *egln3* (prolyl hydroxylase 3) and *epoa* (erythropoietin) in the very same samples. RT-qPCR revealed a 2.1 fold increased expression for *egln3* and 1.5 for *epoa* by 15dpf in *eng*^*−/−*^ fish when compared to siblings. Hypoxia responsive gene expression steadily rose to 6.8 and 14.9 fold increase of *egln3* and *epoa* respectively in *eng*^*−/−*^ in regard to siblings by 30dpf (Fig. 3B). To verify that early difference in *nppa* and *nppb* would reflect heart response to increase workload rather than a direct response to hypoxia (Stockmann et al., 1988), we measured ventricule volume of siblings and *eng*^*−/−*^ at 10, 12 and 15dpf. While this volume was similar between *eng*^*−/−*^ and siblings at both 10 and 12dpf, it appeared significantly increased in mutants at 15dpf (Fig. 3C). Although similar to stress markers in its trend, hypoxia response appeared far more limited in its extent thus strongly suggesting that hypoxia triggered cardiomegaly in *eng*^*−/−*^. Finally, to define how the heart would specifically respond to hypoxic cues in Endoglin deficient fish, we performed FACS analysis to monitor changes in cardiomyocyte abundance. Analysis of cmlc2:GFP^pos^ cardiomyocytes in cell suspensions from whole fish revealed a recurrent increase of GFP^pos^/total cell ratio in 15dpf and older *eng*^*−/−*^ indicating enhanced cardiomyocyte proliferation (Fig. 3D) consistent with a direct proliferative effect of hypoxia on cardiomyocytes (Jopling et al., 2012).

**Fig. 3.**
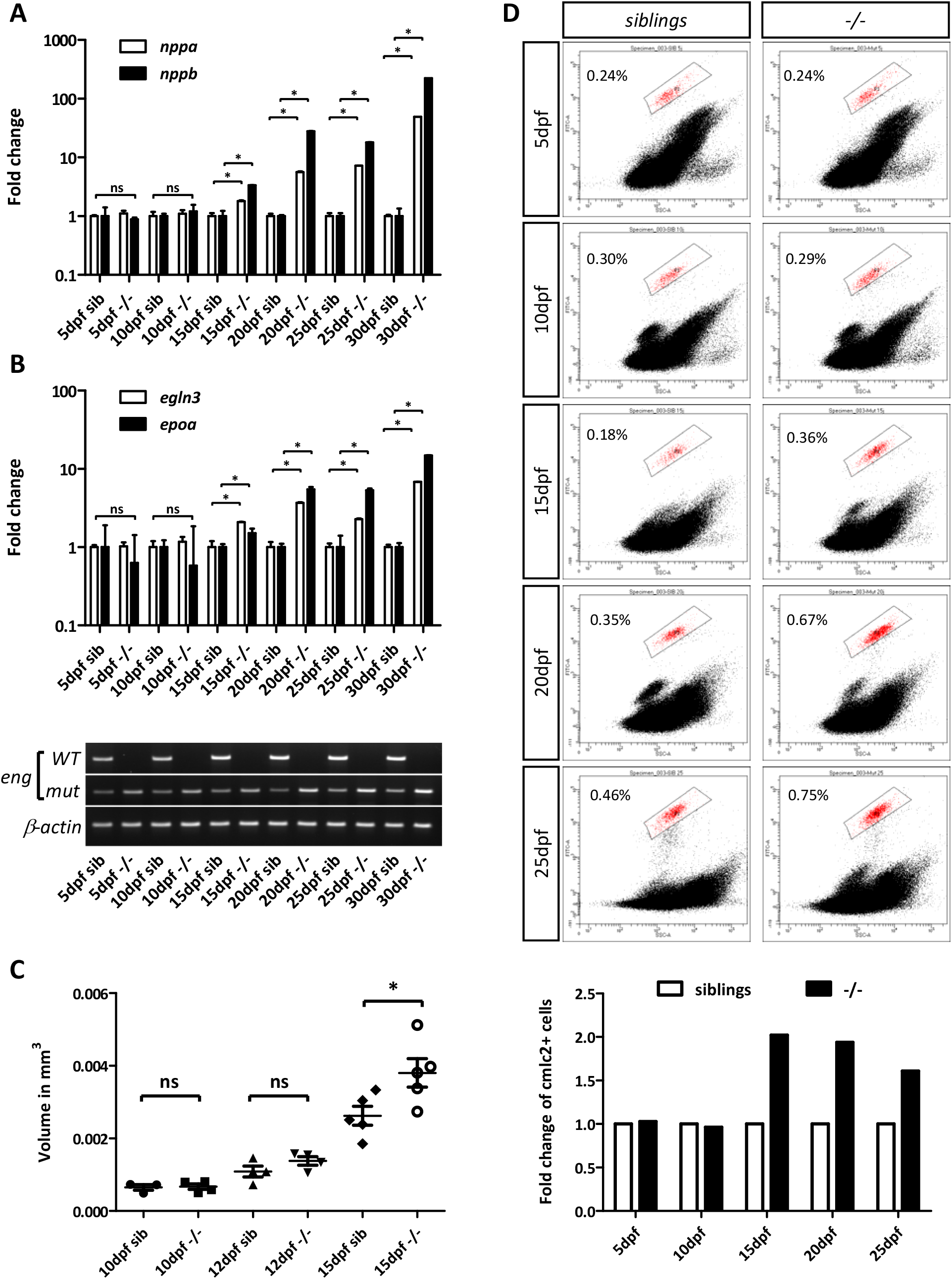
Cardiac stress, ventricule enlargement and cardiomyocyte proliferation correlate with hypoxia in Endoglin deficient fish. (A) Kinetic RT-qPCR analysis of *nppa* and *nppb* (cardiac stress), (B) *egln3*, *epoa* (hypoxia), in 5, 10, 15, 20, 25 and 30 dpf sibling and *eng*^*−/−*^ (genotyping is presented below graphs). Gene of interest expression values were normalized to *rpl13a*. 5dpf=15 fish, 10dpf=15 fish, 15dpf=8 fish, 20dpf=5fish, 25dpf=4 fish, 30dpf=3 fish. Data are mean ± sem of technical replicates. Statistical analysis one-tailed Mann-Whitney test (*nppa* 15dpf *P*=0.05, 20dpf *P*=0.05, 25dpf *P*=0.05, 30dpf *P*=0.05) (*nppb* 15dpf *P*=0.05, 20dpf *P*=0.0383, 25dpf *P*=0.05, 30dpf *P*=0.05), (*egln3* 15dpf *P*=0.05, 20dpf *P*=0.05, 25dpf *P*=0.03831, 30dpf *P*=0.05), (*epoa* 15dpf *P*=0.05, 20dpf *P*= 0.05, 25dpf *P*=0.05, 30dpf *P*=0.05). (C) Analysis of ventricule volume in 10, 12 and 15dpf siblings and *eng*^*−/−*^ in *Tg(cmlc2:GFP)* background. Statistical analysis: one-tailed Mann-Whitney test, (*) *P*=0.0159. (D) Representative Flow cytometry analysis of cardiomyocytes (cmlc2:GFP^pos^) fraction in whole organism cell suspension from 5, 10, 15, 20 ad 25dpf fish. Cell suspensions at 5, 10, 15, 20 and 25dpf were prepared from pool of 26, 18, 9, 7 and 7 sibling or *eng*^*−/−*^, respectively. Bottom graph represents fold change in cmlc2:GFP^pos^ cells in *eng*^*−/−*^ relative to siblings

### Hypochromic Anemia is not the primary trigger of heart failure in Endoglin deficient fish

As anemic mutants such as *riesling* (Liao et al., 2000), *merlot*, *chablis* (Shafizadeh et al., 2002) or *retsina* (Paw et al., 2003) were reported to develop cardiomegaly as the result of cardiac compensation for oxygen transport, we sought to define whether hypochromic anemia observed in *eng*^*−/−*^ fish could account for hypoxia detected by 15pdf. We thus performed a series of o-dianisidine staining in a time course that spanned over the onset of hypoxia and cardiac stress i.e 3, 5, 10 and 15dpf, to evaluate hemoglobin content. No diminished staining was observed in *eng*^*−/−*^ at any given time point when compared with siblings with, inversely, a trend towards enhanced staining in *eng*^*−/−*^ group at 15dpf which would likely reflect increased *epoa* levels at this point (Supplementary Fig. S4). This results indicated that anemia was not the primary cause of hypoxia but a late secondary acquired feature of Endoglin deficiency that would undoubtedly enhance hypoxia.

### Endoglin deficient fish exhibits defects in gill blood vessel development

Since in *eng*^*−/−*^ fish increased *epoa* levels and cardiac compensation failed to correct hypoxia, we hypothesized that chronic hypoxia might reflect respiratory issues. This was supported by bleedings from the gills observed in F_0_ indel carriers. Analysis of histological sections from 30dpf fish revealed that *eng*^*−/−*^ gills were structurally abnormal. Lamellae were unusually short and crooked and supplying blood vessels were markedly enlarged notably in the fourth branchial arch (AA6) (Fig. 4A). We thus compared sibling and *eng*^*−/−*^ gill vascular architecture at 10, 12 and 15dpf in *Tg(kdrl:GFP), Tg(flt-1:Tomato)* transgenic background (Fig. 4B). Although gill vasculature is an arterio-arterial system (Olson, 2002), Flt-1 arterial marker was found mostly associated with afferent branches, whereas *kdrl* promoter appeared evenly active in both afferent and efferent parts of the gill vasculature. As early as 10dpf, differences could already be noticed: vascular loops (afferent/efferent) were shorter in mutants (arrowheads) and the efferent artery of the fourth branchial arch was substantially dilated (asterisk). This differences exacerbated later on and by 15dpf while pan-vascular endothelium *kdrl* reporter expression remained essentially unchanged, we observed a dramatic loss of Flt1 reporter activity in the afferent arterial gill vascular network suggesting that arterial identity might be affected. Finally, to ascertain Endoglin direct role in gill formation, we analyzed endoglin expression by whole-mount *in situ* hybridization on wild-type fish. Endoglin transcripts were detected in developing gills of 10dpf larvae although expression appeared rather faint, it followed branchial arches pattern and was also detected in the paired anterior dorsal aortas. Expression strengthened and spread out by 12 and 15dpf indicating that Endoglin expression was not restricted to major arteries supplying and collecting blood but also in filamental arteries and lamellae (Figure S5A and S5B). Notably, Endoglin expression appeared absent in heart at these stages indicating that heart failure in *eng*^*−/−*^ arose from extrinsic issues. These data showed that, in *eng*^*−/−*^ fish, important alterations of gill vasculature worsening with time took place ahead of hypoxia and suggested that defective gill function resulted in hypoxemia leading to tissue hypoxia.

**Fig. 4.**
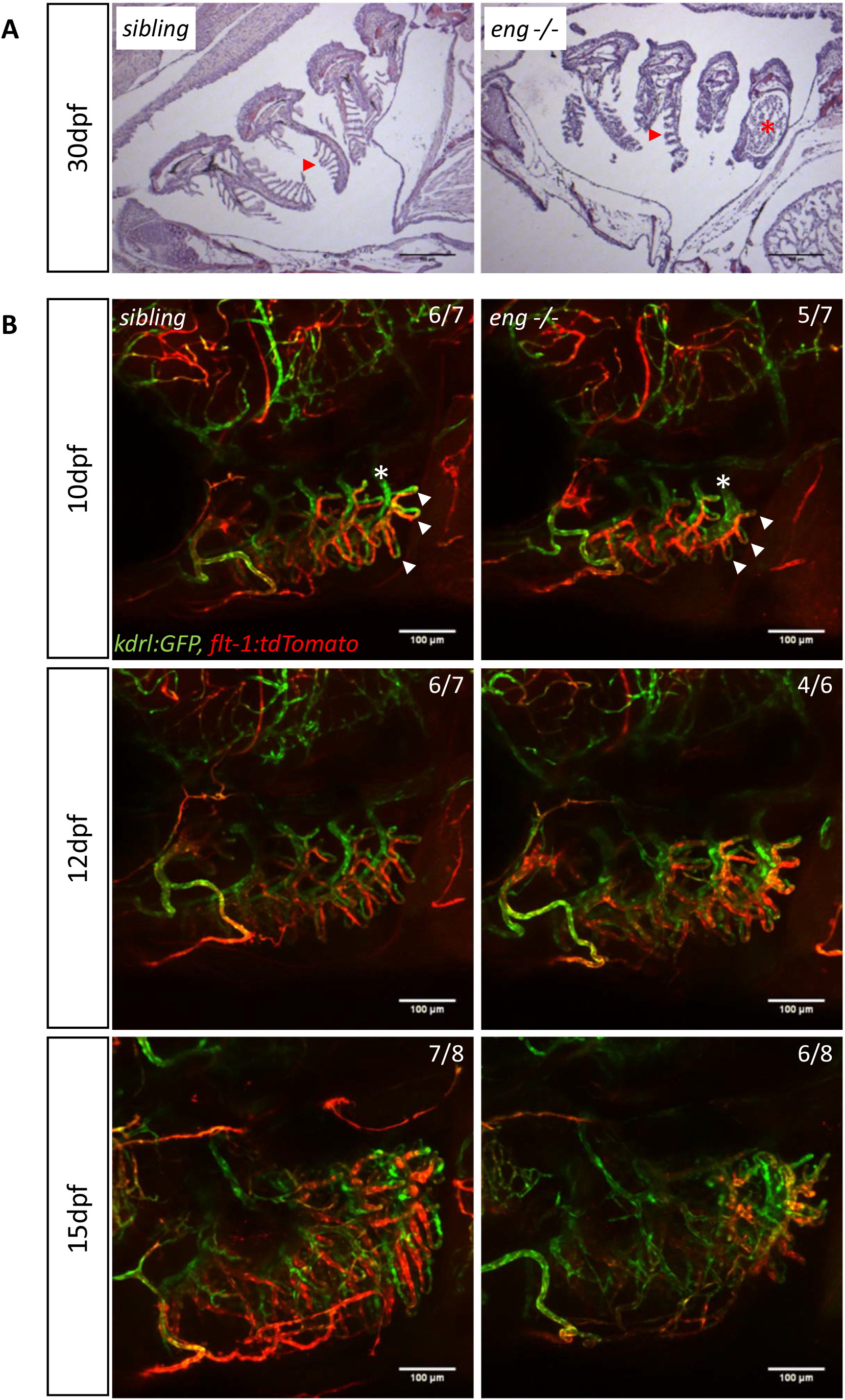
Abnormal branchial vascular development in Endoglin deficient fish underlies structural impairment of gills. (A) Representative histological sections of 30dpf sibling and *eng*^*−/−*^ gills stained with hematoxylin-eosin. Note the poorly developed and organized lamellae (arrowhead) and dramatically enlarged artery (asterisk) on branchial arch 4 (AA6) in *eng*^*−/−*^ fish gills. Bar, 100 μm. (B) Kinetic analysis of gill vascular development in 10, 12 and 15dpf sibling and *eng*^*−/−*^ in Tg(*kdrl:GFP*, *flt1:tdtomato*) background. Note the reduced length of afferent filamental artery (GFP^pos^, dtTomato^High^) and efferent filamental artery (GFP^pos^, dtTomato^low/neg^) (arrowheads) and enlarged efferent branchial artery) (asterisk) as early as 10dpf. Note the overall loss of Flt-1 reporter signal in 15dpf *eng*^*−/−*^. Bar, 100 μm

### Hypoxia-induced erythropoiesis is detrimental to endoglin deficient fish

To define how hypoxia specifically contributed to heart failure in *eng*^*−/−*^ fish, we treated wild-type fish with hemolytic agent phenylhydrazine, a versatile model of hypoxia. To match the onset of hypoxia, phenylhydrazine (phz) was applied every other day from 14dpf up to 75dpf, then fish were allowed to recover. First, we assessed hypoxic and cardiac stress response in 30dpf fish. While hypoxia markers gradually increased with phz concentration and plateaued at 5 μg/ml (Supplementary Fig. S6B upper panels), only phz harshest condition led to a notable increase in cardiac stress markers expression (Supplementary Fig. S6B lower panels), suggesting that *nppa* and *nppb* would be at best weak hypoxia responsive genes in zebrafish. In 5μg/ml phz treated fish, *nppa* reached a 2.33 ±0.16 and *nppb* 6.51 ±0.84 fold activation whereas *egln3* was induced by 3.14 ±0.1 and *epoa* by 2.82 ±0.2 folds. Thus, while hypoxic response in phz treated wild-type and *eng*^*−/−*^ seemed in the same range of magnitude, cardiac stress in phz treated fish appeared overtly marginal (see Fig. 3a, 3b and 5a for comparison). We observed a strict correlation between the extent of heart area enlargement and cardiac stress markers levels but both appeared rather limited compared to *eng*^*−/−*^ at matching age (Supplementary Fig. S6A). Phz treatment had no effect on survival up to this age. Prolonged treatment affected survival in a dose-dependent manner but stayed marginal (Supplementary Fig. S6C). Furthermore, analysis of sex-ratio at 3 month, after one month recovery, did not reveal male gender bias one would expect from a hypoxia-related effect (Supplementary Fig. S6D). Collectively, these results showed that chronic hypoxia *per se* did not result in cardiac issue reminiscent of that observed in *eng*^*−/−*^ fish. We, thus, hypothesized that increased hematocrit/blood viscosity resulting from acute erythropoïeisis and natriuresis (hemoconcentration) could explain these discrepancies. Indeed, 30dpf *eng*^*−/−*^ kidney, fish definitive hematopoïesis organ, displayed increased cellularity, indicative of reactive erythropoïesis, blood smears from 25 and 30dpf *eng*^*−/−*^ revealed high content in erythrocytes, most with immature shape and abnormal staining when compared to siblings and adult surviving *eng*^*−/−*^ showed increased hematocrit when compared to wild-type adult fish (Supplementary Fig. S7A, S7B and S7C respectively). To modulate hematocrit/blood viscosity, we treated *eng*^*−/−*^ and siblings with 0.625, 1.25 and 2.5 μg/ml phz, concentrations that exerted minimal effect on hypoxia and cardiac stress in wild-type fish (Supplementary Fig. S6B). These low phz concentrations showed no overt effect on siblings compared to untreated counterparts. By contrast, phz treated *eng*^*−/−*^, appeared healthier with less prominent cardiomegaly (not shown). We used qPCR to measure both hypoxia and cardiac stress in these different settings (Fig. 5A). Similar to Fig. 3A and 3B, *eng*^*−/−*^ group exhibited a mean 9.3 fold *egln3*, 5.9 fold *epoa*, 31.4 fold *nppa* and 344 fold *nppb*, increase over siblings. On siblings, phz, regardless its concentration, induced modest increases of *egln3*, *epo*a, *nppa* and *nppb* expression over non-treated siblings, while on *eng*^*−/−*^, phz reduced *nppa* and *nppb* expression levels (*nppa* mean fold increase over non-treated siblings group for *eng*^*−/−*^ treated with phz at 0.625, 1.25 and 2.5μg/ml = 18.7, 17.6 and 19.7 respectively) (*nppb* mean fold increase over non-treated siblings for *eng*^*−/−*^ treated with phz at 0.625, 1.25 and 2.5μg/ml = 57.8, 139.6 and 61.9, respectively). Phz also reduced hypoxic response markers expression in *eng*^*−/−*^ (*egln3* mean fold increase over non-treated siblings group for *eng*^*−/−*^ treated with phz at 0.625, 1.25 and 2.5μg/ml = 4.1, 4.5 and 4.8 respectively) (*epoa* mean fold increase over non-treated siblings for *eng*^*−/−*^ treated with phz at 0.625, 1.25 and 2.5μg/ml = 4.2, 4.1 and 6.3, respectively) indicating that heart failure further worsened oxygenation. We, thus, evaluated whether phz treatment would provide long-term benefit on *eng*^*−/−*^ fish health. Fish were treated with either 0.625μg/ml or 1.25μg/ml phz every other day from 14dpf and allowed to recover from 60dpf to 75dpf. As in Fig. 2C, *eng*^*−/−*^ survival was severely impaired compared to siblings with a 16.4% survival at 75dpf (median survival=43 days), while survival of phz treated *eng*^*−/−*^ was markedly enhanced with 41.7% for phz0.625 group and 32.1% for phz1.25 group (median survival of phz0.625 and phz1.25 group equals 61.5 and 60 days, respectively) (Fig. 5B). Further highlighting phz relieving effect on *eng*^*−/−*^ symptoms, treatment allowed higher *eng*^*−/−*^ fractions to reach adulthood and corrected gender bias toward males (Fig. 5C). These results showed that, in *eng*^*−/−*^ fish, hypoxia response failed to work as compensation mechanism but instead brought the pathology to a higher degree.

**Fig. 5.**
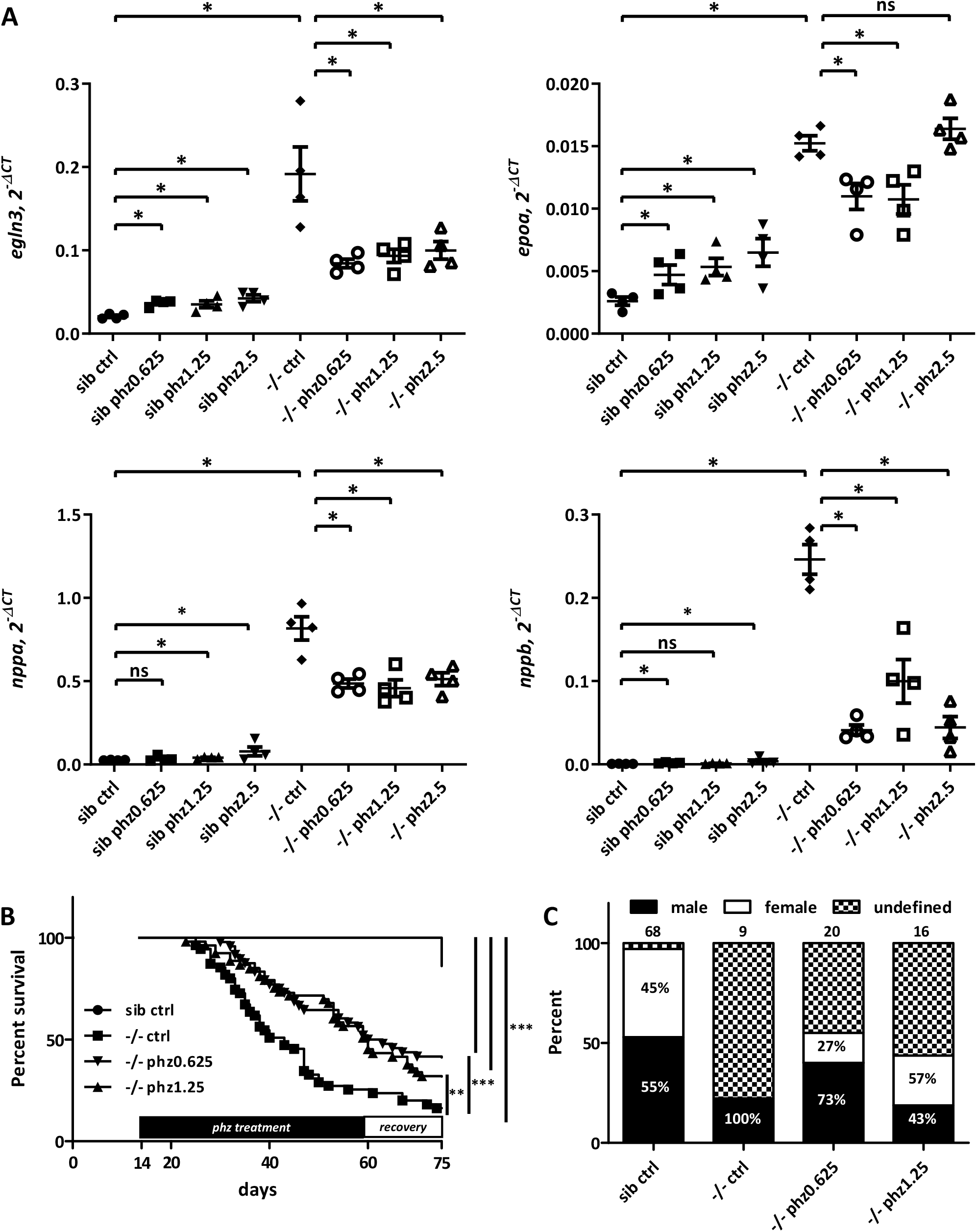
Phenylhydrazine treatment alleviates pathological conditions induced by Endoglin deficiency in zebrafish. (A) RT-qPCR analysis of *egln3*, *epoa* and *nppa* and *nppb* expression in 30dpf siblings (sib) versus *eng*^*−/−*^ fish untreated (ctrl) or treated with 0.625, 1.25 and 2.5μg/ml phenylhydrazine (phz). Target gene expression is represented as 2^−ΔCT^ using rpl13a as reference. Samples are pools of 5 fish. Data are presented as individual sample values and mean ± sem. Statistical analysis one-tailed Mann-Whitney test: *egln3*: sib ctrl vs −/− ctrl *P*= 0.0143, −/− ctrl vs −/− phz0.625 *P*= 0.0143, −/− ctrl vs −/− phz1.25 *P*= 0.0143, −/− ctrl vs −/− phz2.5 *P*= 0.0143; *epoa*: sib ctrl vs −/− ctrl *P*= 0.0143, −/− ctrl vs −/− phz0.625 *P*= 0.0143, −/− ctrl vs −/− phz1.25 *P*= 0.0143; *nppa*: sib ctrl vs −/− ctrl *P*= 0.0143, −/− ctrl vs −/− phz0.625 *P*= 0.0143, −/− ctrl vs −/− phz1.25 *P*= 0.0143, −/− ctrl vs −/− phz2.5 *P*= 0.0143; *nppb*: sib ctrl vs −/− ctrl *P*= 0.0143, −/− ctrl vs −/− phz0.625 *P*= 0.0143, −/− ctrl vs −/− phz1.25 *P*= 0.0143, −/− ctrl vs −/− phz2.5 *P*= 0.0143. (B) Phenylhydrazine treatment enhances *eng*^*−/−*^ survival. Kaplan-Meyer representation of the survival of siblings and *eng*^*−/−*^ non-treated (ctrl) or treated with phenylhydrazine at 0.625 or 1.25 μg/ml. siblings vs −/− ctrl or −/−phz1.25 or −/−phz0.625 *P*<0.0001, −/− vs −/− phz1.25 *P*=0.0039, and −/− vs −/−phz0.625 *P*=0.0009 Log-rank (Mantel-Cox) Test. siblings n=66, −/− n=55, −/−phz1.25 n=53 and −/−phz0.625 n=48. (C) Influence of Phenylhydrazine treatment over sex ratio in 2.5 months plus individuals. Numbers above bars indicate the numbers of surviving fish analyzed. Percent of males and females are indicated inside bars. Of note, fish of undefined gender have been excluded from calculations.

Altogether our results demonstrated that Endoglin deficiency in zebrafish induced heart failure through both direct and indirect effects of hypoxia which stemmed from gill dysfunction. Hypoxia was certainly reinforced by hypochromic anemia and massive hemorrhages from the intestinal tract (Supplementary Fig. S8).

## Discussion

In the present work, we described the generation of a Crispr/Cas9 endoglin zebrafish mutant and the consequences of Endoglin deficiency in post-embryonic stages. We showed that mutant fish massively die from congestive heart failure induced by chronic hypoxia resulting from gill dysfunction. We also found evidence of anemia and observed intestinal hemorrhages that surely further worsen the cardiac pathology. Finally, we showed that health and survival could be strongly enhanced by hemolytic agent phenylhydrazine treatment demonstrating that increased hematocrit/ blood viscosity induced by hypoxia is the main driver of heart failure in Endoglin deficient fish.

Zebrafish endoglin locus expresses two distinct transcripts via the use of an alternate exon introducing an early translation stop resulting in two Endoglin proteins with unique cytosolic sequences. Similar isoforms are also expressed in mammals (Bellón et al., 1993). However, in the latter, Endoglin short isoform is produced by an intron retention mechanism involving Splicing Factor 2 (ASF/SF2) (Blanco and Bernabeu, 2011). Zebrafish Endoglin overall similarity with mammal orthologs is globally low but the cytosolic region of the long isoform is remarkably homologous to that of mammals which interacts with proteins such as GIPC, β-arrestin2 and NOS3 (Lee and Blobe, 2007; Lee et al., 2008; Toporsian et al., 2005). Zebrafish short isoform adds to the wide diversity of sequence of short Endoglin cytosolic domain found in mammals which fluctuates according to the position or the absence of translation stop in retained intron (Blanco and Bernabeu, 2011), and suggests that short isoform cytosolic domain has no protein binding function. Conservation underscores that both Endoglin isoforms should be essential although the short isoform seems absent in reptiles and birds. In mammals, Endoglin short isoform is induced in senescent ECs and transgenic mice overexpressing this isoform are hypertensive and insensitive to NO synthesis inhibitor L-NAME (Blanco et al., 2008). A soluble form (S-Endoglin) is also produced by cleavage of the ectodomain by MMPs and can exert paracrine or remote activity using exosomes as vehicle (Ermini et al., 2017). High S-Endoglin levels correlate with preeclampsia and induce hypertension by inhibiting TGF-β-NOS axis (Venkatesha et al., 2006). Further work will be necessary to establish whether S-Endoglin is actually produced in zebrafish but sequence analysis indicates that ectodomain Glu-Leu residues essential for MMP14 cleavage in mammals, are conserved (Hawinkels et al., 2010).

In zebrafish, by contrast with mammals, Endoglin participation in ALK1 signaling seems highly questionable. Expression patterns marginally overlap during development and differences in lethality timing between ALK1 and Endoglin mutants (7-10 vs 30dpf, respectively) rather argue against a function on a shared pathway(Roman et al., 2002). Moreover, ALK1 mutants exhibit enlarged cranial blood vessels due to impaired EC ability to migrate against blood flow (Corti et al., 2011; Rochon et al., 2016; Roman et al., 2002). These defects are phenocopied by ALK1 or both bmp10 and bmp10-like morphants demonstrating that BMP10s signal through ALK1 to control cranial blood vessel caliber (Laux et al., 2013). This phenotype is, by contrast, absent in Endoglin mutants and morphants (not shown) indicating that Endoglin is dispensable for BMP10-ALK1 signaling in this respect. Conversely, phenotypes of Endoglin mutant (this study and previous report (Sugden et al., 2017)) i.e accelerated blood flow in a simple DA/PCV vascular loop is not described in ALK1 deficient zebrafish embryos indicating that Endoglin contribution to ALK1 pathway is not restricted to a specific vascular bed either. ALK5 seems neither an alternate candidate receptor as both alk5a and alk5b are overtly absent in axial blood vessels by the time Endoglin deficiency induces changes in blood flow pattern (Park et al., 2008). Recent data, in mammals, suggest that Endoglin could act outside of TGFβ/BMP signaling. Endoglin was indeed found to interact with VEGF-R2 and promoted its signaling in response to VEGF (Tian et al., 2018). Further investigation will definitely be required to define whether, in zebrafish, phenotypes induced by Endoglin deficiency could be related to this specific signaling. Interestingly, ALK1 conditional knockout in adult mice results in high output heart failure and anemia (Morine et al., 2017). EC-specific endoglin knock-out in adult mice, also results in high-output heart failure induced by AVM formation in the pubic symphysis as the result of exacerbated VEGF-R2 receptor signaling in Endoglin deficient condition (Tual-Chalot et al., 2020). Thus, ALK1-Endoglin interaction might be required at later stages in zebrafish. ALK1 mutant early lethality precludes analysis of its function in gill vascular development. But, it will be interesting to assess whether constitutively active ALK1 or soluble VEGF-R1 would rescue heart issue in Endoglin deficient zebrafish.

Our results demonstrate that zebrafish Endoglin deficiency elicits a chronic hypoxic response. In normoxia, HIFs (HIF-1α/HIF-2α) are hydroxylated on specific proline by prolyl hydroxylases (PHDs) which use O_2_, 2-oxoglutarate, ascorbate and ferrous iron ions (Fe^2+^) as cofactors. Hydroxylated HIFs are recognized by the Von Hippel Lindau protein (pVHL) component of E3 ubiquitin ligase complex leading to HIFs polyubiquitylation and ultimately HIFs degradation by the proteasome. Conversely, hypoxia hampers PHD activity leading to HIFs stabilization, nuclear translocation and transcription factor complex formation with HIF-1β/ARNT and regulation of specific sets of genes (Ivan and Kaelin, 2017). Iron deficiency leads eventually to anemia due to the essential contribution of iron to the haem complex essential to hemoglobin for oxygen transport. Iron free diet represses iron reabsorption inhibitor Hepcidin by a HIF-dependent transcriptional repression mechanism (Peyssonnaux et al., 2007). In keeping with this, hypoxia mimetics desferrioxamine and cobalt or nickel chloride all inhibit Prolyl hydroxylases (PHD) by interfering with ferrous iron (Muñoz-Sánchez and Chánez-Cárdenas, 2019). Endoglin deficient fish exhibit features highly reminiscent of vhl zebrafish mutants which model Cuvash syndrome. These mutations abolish vhl ability to associate with HIF factors and lead to the constitutive activation of HIF-dependent pathways in normoxia. Vhl mutants exhibit exacerbated erythropoiesis in response to Epo overexpression which results in a form of polycythemia with immature and hypochromic characteristics. Mutant fish also exhibit a hyperventilation phenotype which stems from the direct action of Epo on respiratory neurons and develop congestive heart failure, edema and eventually die by 11dpf (van Rooijen et al., 2009). In mouse, liver-specific deletion of VHL results in HIF-mediated Hepcidin repression and Epo induction leading to excessive (polycythemia) microcytic and hypochromic erythropoiesis. Thus, despite converging mechanisms set to mobilize iron and red blood cell production, conditions of chronic HIF stabilization end up invariably in an imbalance between Epo and iron levels (Peyssonnaux et al., 2007).

Chronic hypoxia induces cardiomegaly in both fish and mammals but unlike mammals where cardiomyocytes undergo size increment (hypertrophy), fish cardiomyocytes are able to re-enter cell cycle after dedifferentiation (Jopling et al., 2010). Our data, although indirectly, show that cardiomyocyte proliferation is indeed enhanced in Endoglin deficient fish in a timetable that matches hypoxia. Recent reports have shed light as to how HIF activation induces cardiomegaly through cardiomyocyte hypertrophy and cardiac remodeling : Hypoxia Inducible Mitogen Factor (HIMF), a non-canonical ligand of Calcium Sensing Receptor (CaSR) controls *in vitro* and *in vivo* cardiomyocyte hypertrophy by activating HIF, CaN-NFAT and MAPKs pathways (Kumar et al., 2018; Zeng et al., 2017). HIMF potential orthologs while present in the genome of some fish i.e. *Erpetoichthys calabaricus*, are, to date, absent in zebrafish databanks.

By contrast with vhl, hypoxic response in endoglin mutants is gradually acquired thus indicating that Endoglin does not control HIF activation. Instead, our data strongly argue in favor of a faulty respiratory system. Fish extract water dissolved oxygen through their gills, a highly complex respiratory organ composed of multiple functional units called lamellae. Bridging afferent and efferent circulation, lamellae are thin flat vascular sinusoids of endothelial and non–endothelial pillar cells ensheathed by a layer of pavement epithelial cells (Olson, 2002). In zebrafish, gills lamellae were found to form around 12-14dpf to reach their definitive adult morphology by about 4 weeks (Rombough and Drader, 2009). They develop on a scaffold of vascular loops composed of afferent and efferent parts linking ventral aorta to lateral and dorsal aorta through branchial arches. Since structural and molecular alterations of the gill vasculature are observed in Endoglin deficient zebrafish before or at the very onset of lamellae formation, these defects would likely directly hamper lamellae development and gill function. Gill dysfunction will result in hypoxemia leading to poor tissue oxygenation triggering hypoxic response. It is, thus, not surprising that mutant fish will start to decline by 30dpf which corresponds to the time when gill structure reaches its fully operative shape in normal zebrafish and that variability in gill dysfunction allows fish to survive longer periods, not excluding, though, that blood vessel defects affecting other organs resulting in inappropriate perfusion would reinforce hypoxia.

Treating Endoglin deficient fish with hemolytic agent phenylhydrazine has a strong impact on fish health. Enhanced survival clearly demonstrates that it protects from heart failure. This beneficial effect of phenylhydrazine, a drug historically used to cure polycythemia vera (Long, 1926), suggests that a form a polycythemia is taking place in Endoglin deficient fish that puts the heart at stake. Interestingly, in HHT, polycythemia is a complication of pulmonary AVM which, by mixing arterial and venous blood (right-to-left shunting), results in hypoxemia eventually triggering erythropoiesis (for review (Cottin et al., 2007; McDonald et al., 2011)). Thus, despite structural difference between fish and human cardiorespiratory systems, Endoglin deficiency appears to results in similar pathological situations.

Zebrafish has become a choice organism to perform functional screens of bioactive molecules of therapeutic interest. Hence, by reproducing important features of HHT, both heterozygous and homozygous Endoglin mutant fish will give us the opportunity to address the 2^nd^ hit issue by screening for drugs able to induce symptoms at a higher frequency, while on the other hand homozygotes will serve to identify molecules able to rescue both embryonic and post-embryonic phenotypes.

## Experimental procedures

### Zebrafish lines, maintenance and ethics

All zebrafish (*Danio rerio*) lines were produced or maintained in AB genetic background. Transgenic lines used in this report are: *Tg(myl7:EGFP) here referred to as Tg(cmlc2:EGFP)*, *Tg(−0.8flt1:RFP) here referred to as Tg(flt1:Tomato), Tg(kdrl:GFP), Tg(−8.1gata1a:mRFP) here referred to as gata1:RFP*. Embryos were staged according to Kimmel et al. (Kimmel et al., 1995) and blood vessels nomenclature refers to Isogai et al. (Isogai et al., 2001). All experiments were performed in accordance with the 2010/63/EU Directive and the ARRIVE guidelines (Kilkenny et al., 2010). Procedures used in this study have been approved by Ethical Committee (#036) as part of an authorized project registered by the French Ministère de l’Enseignement Supérieur de la Recherche et de l’Innovation under APAFIS number #23822-2020052717421507 v3 (E. Lelièvre). Animals were housed in a zebrafish facility registered under Agreement number #A3417237.

### Total RNA extraction, semi-quantitative and qRT-PCR analysis

Total RNA was extracted from staged embryos, sibling and *eng*^*−/−*^ fish at specific ages as pool using Nucleospin RNA kit (Macherey-Nagel) or NucleoZOL (Macherey-Nagel) following manufacturer’s instructions. Equal amounts (500 ng or 1 μg) of total RNA were retrotranscribed using High Capacity cDNA Reverse Transcription kit (Applied Biosystems) following manufacturer’s instruction. Endoglin variants were amplified by semi-quantitative PCR using Phusion Hot Start II High-Fidelity DNA polymerase (Thermo Scientific) or HotGoldStar PCR mix (Eurogentec) using 50 ng total RNA equivalent RT reaction and Endoglin EngExon12fwd 5’-CTGGGCATAGCGTTCGGAGGATT-3’, EngExon10fwd 5’-CCTGGGACCTCGAATGTGCTGTAA-3’ EngExon15rev 5’-CATGCTGCTGGTGGGTGTGCT-3’ specific primers and Housekeeping gene beta-actin bactinfwd 5’-CCTGGAGAAGAGCTATGAGCTG-3’, bactinrev 5’-ATGGGCCAGACTCATCGTACTC-3’ primers as control. When necessary, the genotype of samples was verified by RT-PCR using EngWTfwd, EngMutfwd (see sequence below) and dEngExon5rev 5’-CGTTGGTGACGGATGTGACT-3’ according to the semi-quantitative PCR procedure described above. qPCR reactions were carried out using 1.25 ng total RNA equivalent RT reaction in sensiFAST SYBR No-Rox mix (Bioline) in presence of 600 nM of each primer. Reactions were assembled in triplicates in 384-well plates using Labcyte Echo 525 Liquid Handler and PCR reaction was performed using Roche LightCycler 480 available at MGX - High throughput qPCR facility. PCR reactions were conducted using following primers : dreegln3fwd 5’-TGGGAAAAAGCATTCGTGCG-3’, dreegln3rev 5’-CGGCCATCAGCATTAGGGTT-3’, dreepoafwd 5’-CCATTACGCCCCATCTGTGA-3’, dreepoarev 5’-GTGACGTTCGTTGCAATGCT-3’, drenppafwd 5’-GACACAGCTCTGACAGCAACA-3’, drenpparev 5’-TCTACGGCTCTCTCTGATGCC-3’, drenppbfwd 5’-TGTTTCGGGAGCAAACTGGA-3’, drenppbrev 5’-GTTCTTCTTGGGACCTGAGC-3’, drerpl13afwd 5’-CGCTATTGTGGCCAAGCAAG-3’, drerpl13arev 5’-TCTTGCGGAGGAAAGCCAAA-3’. dreengfwd 5’-AGACGGAGAACGGGACAGAA-3’, dreengrev 5’-TCACCACAGACTTGTTCGCC-3’.

### 5’ and 3’ Rapid Amplification of cDNA End (RACE) experiments

For endoglin 5’ RACE we used a template-switch strategy based on a previously described procedure (Pinto and Lindblad, 2010). Briefly, 36hpf total RNA (1 μg) were retrotranscribed from either 5’-CATGCTGCTGGTGGGTGTGCT-3’ or 5’-CCATAAAGCACCGGTGAGCAGAA-3’ endoglin specific reverse oligonucleotides (500 nM) using RevertAid H minus reverse Transcriptase (Thermo Scientific) 10 U/μl in 1X RevertAid H minus Buffer supplemented with 2 U/μl RNAse OUT (Thermo Scientific), 1 mM dNTPs (Thermo Scientific), 2 mM MgCl_2_ at 50°C for 1 hour. The template-switch reaction was then conducted in presence of Template Switch oligonucleotide (1 μM) and 3 mM MnCl_2_ and 4 U/μl RevertAid H minus reverse transcriptase for 90 minutes at 42°C. Reverse transcriptase was finally inactivated by a 10 min at 70°C step. Then using Template Switch RTs (50 ng total RNA equivalent) as template together with shortened U_sense oligonucleotide 5’-GTCGCACGGTCCATCGCAG-3’ and endoglin-specific oligonucleotides 5’-CCATAAAGCACCGGTGAGCAGAA-3’ and 5’-GTTTATCCTTTTGACGCCGCAGAG-3’ as primers, PCR reactions were performed using Phusion Hot Start II High-Fidelity DNA polymerase (Thermo Scientific) following manufacturer’s instructions. Endoglin 3’RACE was performed using GeneRacer kit (Thermo Scientific) following manufacturer’s recommendations. Briefly, 36hpf total RNA (1 μg) were used in RT reactions containing 2.5 μM of GeneRacer Oligo dT primer and 10 U/μl of SuperScript III RT. PCR reactions were then conducted as described above using endoglin-specific forward primer 5’-CCTGGGACCTCGAATGTGCTGTAA-3’ and GeneRacer 3’ primer. One percent of whole PCR was then used as template in nested PCR reactions using endoglin-specific forward primer 5’-CTGGGCATAGCGTTCGGAGGATT-3’ and GeneRacer 3’ Nested primer. PCR products from 5’ and 3’RACE were then cloned blunt into pBSSK(−) vector and sequenced on both strands using T3 and T7 primers.

### Whole-mount in situ hybridization

Whole-mount *in situ* hybridization was performed essentially as previously described (Thisse and Thisse, 2008). Briefly, wild type AB fish embryos or larvae were fixed overnight at 4°C in 4% PFA, pH 9.5. Digoxygenin-labeled Endoglin antisense riboprobe was synthesized from Endoglin partial cDNA obtained from 5’ RACE (nucleotide 1 to 1454) cloned into pBSSK(−) vector. Samples were pre-hybridized overnight at 65°C and hybridization was performed overnight at 65°C in presence of 0.4 ng/nl endoglin antisense probe. Probe was detected using alkaline phosphatase conjugated anti-digoxygenine Fab fragments (Roche) and NBT/BCIP reagent mix (Roche).

### CRISPR/Cas9 generation of endoglin mutant and genotyping

One nanoliter of a mixture composed of endoglin targeting gRNA S. pyCas9-3NLS recombinant protein in 20 mM HEPES pH 7.5, 150 mM KCl was injected into 4- to 16-cell-stage wild type AB embryos. Cas9 recombinant protein and gRNAs were purchased from TACGENE. Fish were raised to adulthood and indels carriers were screened by T7 Endonuclease I (T7EI) assay following PCR amplification from crude genomic DNA. T7EI positive samples were sequenced and analyzed using TIDE software (https://tide.nki.nl/). Selected founder F_0_ candidates were crossed with wild type AB and progeny (F_1_) raised to adulthood. Indels carriers were identified using T7EI assay and direct sequencing of PCR product. A single endoglin mutant line was maintained for further analysis. F_1_, F_2_ or F_3_ mutation carriers were then crossed with transgenic reporter lines maintained in AB background. Throughout this study Endoglin mutation was maintained in a heterozygous status to prevent the potential selection of individuals able to cope with Endoglin deficiency. Routine genotyping was performed on crude genomic DNA prepared from caudal fin clips or in the case of survival experiments pieces from dead larvae or juveniles using an allele specific PCR strategy with engWTfwd 5’-ACAGACGAATCTACAGCCGACAT-3’, engMutfwd 5’-AGAACAGACGAATCTACAGCCGAA-3’ and engintron2rev 5’-AGCATGTTTTAACAAGACGGCAG-3’ primers and HotGoldStar PCR mix (Eurogentec).

### Survival

Dead fish were collected for genotyping to confirm sibling vs *eng*^*−/−*^ status and discriminate wild-type from heterozygous. Kaplan-Meier survival plots and Log-Rank (Mantel-Cox) statistical analysis were generated using Prism GraphPad software.

### Imaging of blood vessel perfusion

Seventy-two hours post-fertilization (72hpf) GFP^pos^ and dsRed^pos^ sibling and *eng*^*−/−*^ from crosses between *eng*^*+−/−*^, *Tg(kdrl:GFP*) and *eng*^*+−/*^, *Tg(gata1:dsRed)* fish were anesthetized mounted in 0.7% low melt agarose, MS-322 (160 μg/ml) and imaged on Zeiss AXIO Zoom.V16 mounted with Zeiss AxioCam MRm with Zeiss HXP 200C illuminator set on minimal power using fixed exposure parameters (GFP 1 sec and CY3 1.5 sec) to obtain red blood cells traces revealing perfusion extent. Similar post-acquisition treatments were applied to images.

### Tissue clearing and deep imaging of whole zebrafish

To get access to *in situ* dimensions of siblings and *eng*^*−/−*^ heart ventricule at 10, 12 and 15dpf, Clutches from crosses between *eng*^*+/−*^,*Tg(cmlc2:GFP)* and *eng*^*+/−*^ were screen at 24dpf to sort out GFP^pos^ individuals and at 72dpf to discriminate siblings from *eng*^*−/−*^. Fish were raised as described earlier and collected at indicated times by excess of ethyl 3-aminobenzoate methanesulfonate (MS222) (320 μg/ml) and fixed overnight in 4% PFA. Whole zebrafish were depigmented and labeled according to previously published protocol (Frétaud et al., 2021) using Rabbit anti-mCherry (Rockland, 600-401-P16) and Chicken anti-GFP (ThermoFisher, A10262) prior to Alexa594 goat anti-rabbit (ThermoFisher, A11012) and Alexa488 goat anti-chicken (ThermoFisher, A11039). Before imaging, larvae were cleared by incubation in RIMS (Yang et al., 2014) overnight at RT. Larvae were mounted under #1 coverslips in RIMS supplemented with 0.8 % low gelling agarose. Images were acquired with a Leica SP8 confocal microscope using a HCX IRAPO L 25X/0,95NA water immersion objective (#11506340, Leica microsystems). Ventricule volume is estimated by the simplified ellipsoid volume calculation formula V=0.523(width in mm)^2^(length in mm) (Hoage et al., 2012).

### Confocal imaging

GFP/Tomato positive embryos from crosses between *eng*^*+/−*^, *Tg(kdrl:GFP)* and *eng*^*+/−*^, *Tg(flt1:Tomato)* were screened at 72hpf to discriminate siblings from *eng*^*−/−*^. Fish were raised as described earlier and collected at indicated times by excess of MS222 (320 μg/ml) and fixed overnight in 4% PFA. Fixed zebrafish were mounted in 0.7% low-melt agarose in Fluorodish cover-glass bottom culture dish (WPI) and imaged on a Zeiss LSM510 confocal microscope. Projections of Z-stack were performed using Fiji software. Similar post-acquisition treatments were applied to images.

### Flow cytometry

GFP^pos^ siblings and *eng*^*−/−*^ fish from crosses between *eng*^*+−/−*^ and *eng*^*+−/−*^, *Tg(cmlc2:EGFP)* fish were sorted at 3dpf and raised along as described above. At 5, 10, 15, 20 and 25dpf fish were collected as pool of 26, 18, 9, 7 and 7 fish respectively and dissociated in 0.25% Trypsin-EDTA (Gibco, ThermoFisher Scientific) and 8 mg/ml Collagenase from *Clostridum histolyticum* (Sigma-Aldrich) according to previously published procedure (Bresciani et al., 2018). Dissociated cells were fixed overnight in 4% PFA at 4°C, then rinsed thrice in PBS and store at 4°C in PBS, 0.05% sodium azide until analysis. 250000 cells were analyzed on a FACSCanto cytofluorimeter (Becton Dickinson) at each indicated time point and genotype to evaluate cardiomyocytes (GFP^pos^) representation in whole fish cell suspensions. FACs profiles were validated using cardiomyocytes enriched cell suspensions prepared from GFP^pos^ isolated *Tg(cmlc2:EGFP)* hearts.

### Phenylhydrazine treatment

Wild-type zebrafish (about 30 fish per batch) were bathed every other day from 14dpf with 1.25, 2.5 and 5 μg/ml phenylhydrazine in fish water prepared from a freshly made 5 mg/ml stock solution of phenylhydrazine hydrochloride (Sigma-Aldrich). Non-treated fish served as control. Bathing volume was gradually increased to accommodate with fish growth (100 to 300 ml). Fish were treated for 30 min and then allowed to recover in fresh fish water for about 1 hour and finally replaced into housing tanks and fed. Fish were subjected to this regimen up to 75dpf before treatment was definitively stopped to let fish recover and regain gender-specific hallmarks to allow for accurate sex ratio determination. To assess hypoxia and cardiac stress in fish under phenylhydrazine treatment, 4 batch of 5 fish of each condition were collected at 29dpf (treatment-free day) and euthanized by excess of MS222 (320 μg/ml) for total RNA extraction performed as described above. Non-treated wild-type fish were used as control. Phenylhydrazine treatment of *eng*^*−/−*^ and siblings was performed as for wild-type (see above) but phenyhydrazine concentrations were downscaled to 0.625, 1.25 and 2.5 μg/ml. Non-treated Siblings and *eng*^*−/−*^ were used as control. At 29dpf (treatment-free day), each condition was divided into 4 samples of 4-5 fish/sample and used for total RNA extraction. For phenylhydrazine effect on *eng*^*−/−*^ survival, siblings and *eng*^*−/−*^ fish were treated as described above using 0.625 and 1.25 mg/ml phenylhydrazine up to 2 months and were left to recover for 15 days to allow for gender determination.

### Histology

Fish were euthanized by excess of MS222 (320 μg/ml) and fixed overnight in 4% PFA. Samples were decalcified using 0.35 M EDTA. Paraffin embedding was performed by RHEM Histology Facility. Sagittal sections (7 μm) were dewaxed and stained by hematoxylin-eosin. WISH samples were post-fixed overnight in 4% PFA. Paraffin embedding was performed by RHEM. Sagittal and transversal sections (7 μm) were dewaxed and counterstained with Nuclear Fast Red.

### Hemoglobin staining, blood smears and hematocrit measurement

Hemoglobin content in 3, 5 10 and 15dpf siblings and *eng*^*−/−*^ fish was assessed using o-dianisidine reagent according to previously published procedure (Leet et al., 2014). Briefly, embryos and larvae were stained with 0.6 mg/ml o-dianisidine (Alfa Aesar) in 10 mM sodium acetate pH4.5, 0.65% H_2_0_2_ and 40% (v/v) ethanol for 30 min at room temperature. Fish were then wash with PBS and fixed overnight in 4% PFA. The next day, fish were washed with PBS and soaked for 30 min in 0.8% KOH, 0.9% H_2_0_2_ and 0.1% Tween-20 to remove pigments. After washes with PBS, fish were scored for hemoglobin staining and pictures of representative fish were taken after mounting in 0.7% low-melt agarose. For blood smears, fish were anaesthetized using MS222 (160μg/ml) and blood samples obtained from cardiac puncture were smeared on microscope slides. Wright-Giemsa staining was performed following manufacturer’s instructions (Sigma-Aldrich). For hematocrit measurement, wild-type and *eng*^*−/−*^ adult fish were anaesthetized as described above and blood samples were obtained by cardiac puncture using 18 μl heparinized capillary tubes (Hirschmann). Samples were processed using Hematocrit 24 centrifuge (Hettich) and hematocrit was assessed.

## Acknowledgements

Authors thank Mireille Rossel, Nicolas Cubedo, Philippe Clair and Stéphane Delbecq for sharing their expertise.

## Data availability

The data underlying this article are available in the article and online supplementary material. Sequence information from endoglin 5’ and 3’ RACE will be made available upon request.

## Author contributions

EL. CB. YB. MF. CL. and CJ Performed experiments and/or analyzed data. EL with input from CJ and KK conceived and designed the study. EL wrote the manuscript.

