## supplemental data for "Endoglin deficiency elicits hypoxia-driven congestive heart failure in zebrafish"

**Legends to supplementary figures**

**Supplementary Fig. S1. Decreased amounts of endoglin transcripts in *eng*$^{-/-}$ suggest nonsense mRNA decay of mutant transcripts.** Quantitative RT-PCR analysis of all-endoglin (A), wild-type endoglin (B) and mutant endoglin (C) mRNA expression in 72hpf wild-type (WT), siblings (sib) and *eng*$^{-/-}$ (-/-) embryos. Target gene expression is represented as $2^{-\Delta CT}$ using rpl13a as reference. Samples (5 for each genotype) are pools of 17 to 23 embryos. nd, not detected. Statistical analysis: one-tailed unpaired *t*-test; WT vs sib *P*=0.0984, WT vs -/- *P*=0.0012, sib vs -/- *P*=0.0093.

**Supplementary Fig. S2. Overall appearance of 3-month-old adult size *eng*$^{+/+}$, asymptomatic and symptomatic *eng*$^{+/-}$ and *eng*$^{-/-}$ fish**. Note the overall reddish color of symptomatic fish and dilated blood vessels at the root of caudal, pectoral and anal fins. Arrow points to enlarged cardiac area. Pictures were taken from alive fish directly in their housing tanks therefore fish size is not representative

**Supplementary Fig. S3. Phenotypic details of symptomatic *eng*$^{+/-}$.** Representative lateral (left panels) and ventral (right panels) pictures of adult (6 months) *eng*$^{+/+}$, asymptomatic and symptomatic *eng*$^{+/-}$ and *eng*$^{-/-}$ (females) showing the presence of dilated blood vessels reminiscent of telangiectasias. Note the cardiomegaly in *eng*$^{-/-}$ and characteristic hydropsy symptoms such as bulging eyes and pinecone-like scales, sign of multiple organ failure only observed secondary to heart failure. Bar, 1mm

**Supplementary Fig. S4. Anemia is not responsible for hypoxia in Endoglin deficient larvae.** Hemoglobin assessment in siblings versus *eng*$^{-/-}$ does not reveal early anemia in *eng*$^{-/-}$. Representative Whole-mount o-dianisidine staining of 3, 5, 10 and 15dpf *eng*$^{-/-}$ and siblings. Numbers in upper right corner of pictures indicate the number of fish with similar pattern out of total number analyzed. Bar, 500µm

**Supplementary Fig. S5. Endoglin expression in developing gills.** (A) Whole-mount in situ hybridization using Endoglin antisense riboprobe on 10, 12 and 15dpf wild-type zebrafish larvae. Upper panels, lateral views; lower panels, ventral views. Bar, 100µm. Asterisk indicate heart location. (B) Representative histological sections of WISH using endoglin antisense probe in 15dpf zebrafish larvae showing endoglin expression in developing gills. Sagittal section (left) transversal section (right) (b,brain; ea-lda, epibranchial arteries-lateral dorsal aorta; g, gills k, kidney; l, liver, va, ventral aorta). Bar, 100µm

**Supplementary Fig. S6. Phenylhydrazine-induced hypoxia fails to mimic Endoglin deficiency heart failure condition.** (A) Representative pictures of one-month-old wild-type fish non-treated (ctrl) or treated with 5 µg/ml phenylhydrazine (phz). Note the overt paleness of gills area (asterisk) and slightly enlarged heart region (arrowhead). Bar, 1mm. (B) RT-qPCR analysis of egln3, epoa (hypoxia responsive gene) and nppa and nppb (cardiac stress responsive gene) expression in one-month-old wild-type fish non-treated (ctrl) or treated with 1.25, 2.5 and 5µg/ml phenylhydrazine. Target gene expression is represented as $2^{-\Delta CT}$ using rpl13a as reference. Samples (4 for each condition) are pools of 5 fish. Data are presented as individual sample values and mean ± sem. Statistical analysis: one-tailed Mann-Whitney test; *egln3* ctrl vs phz1.25 and phz2.5 $P$=0.0097, ctrl vs phz5 $P$=0.0060, phz1.25 vs phz2.5 $P$=0.0143, phz1.25 vs ph5 $P$=0.0079; *epoa* ctrl vs phz1.25 $P$=0.0286, ctrl vs phz2.5 and phz5 $P$=0.0143, phz1.25 vs phz2.5 and ph5 $P$=0.0143; *nppa* ctrl vs phz2.5 and phz5 $P$=0.0143, phz1.25 vs ph5 $P$=0.0143; *nppb* ctrl vs phz2.5 and phz5 $P$=0.0143, phz1.25 vs ph5 $P$=0.0143. ns $P$>0.05. (C) Analysis of phenylhydrazine treatment effect over fish survival. Kaplan-Meyer representation of the survival of non-treated (ctrl) and 1.25, 2.5 and 5 µg/ml phz treated wild-type fish. Note the dose dependent effect of phz concentration over survival but absence of deleterious effect at early juvenile stage. ctrl vs phz1.25 $P$=0.0114, ctrl vs phz2.5 $P$=0.0051, ctrl vs phz5 $P$<0.0001, phz1.25 vs phz5 $P$=0.0113, phz2.5 vs phz5 $P$=0.0310 and phz1.25 vs phz2.5 $P$=0.71219 (ns) Log-rank (Mantel-Cox) Test. (D) Analysis of phz treatment influence over sex ratio in 3-month-old individuals. Note the absence of gender bias at any phenylhydrazine concentration.

**Supplementary Fig. S7. Hematological features of *eng*$^{-/}$ fish.** (A) Representative pictures of hematoxillin-eosin stained histological sections from siblings versus *eng*$^{-/-}$ fish at 30dpf. Note kidney hypercellularity indicative of reactive erythropoiesis. Bar, 100µm. (B) Representative pictures of Wright-Giemsa stained blood smears from sibling versus *eng*$^{-/-}$ fish at 25 and 30dpf. Note the marked difference of erythrocyte shape and staining between siblings and *eng*$^{-/-}$. Bar, 20µm. (C) Adult (6 month or older) *eng*$^{-/-}$ exihibit increased hematocrit compared to wild-type fish. Mean hematocrit of wild-type (n=13) 41.96 ± 1.798 % vs *eng*$^{-/-}$ (n=13) 50.75 ± 1.746 %. Statistical analysis: one-tailed unpaired *t*-test; $P$ =0.0009.

**Supplementary Fig. S8. Intestinal hemorrhages in adult *eng*$^{-/-}$ fish.** (A) Wright-Giemsa stained smear of collected clot from hemorrhage in 5 month old fish. The smear (only part of it) is shown as an indication of blood loss extent. Bar, 100µm. (B) Fresh clot from intestinal hemorrhage from 2-month-old *eng*$^{-/-}$ fish. Bar, 1mm. Inset, close-up picture. Of note, the size of the sample indicative of massive blood loss. Clot rapidly turns pale since erythrocytes tends to release their content once exposed to tank water. Fish died 4 days after hemorrhagic episode.

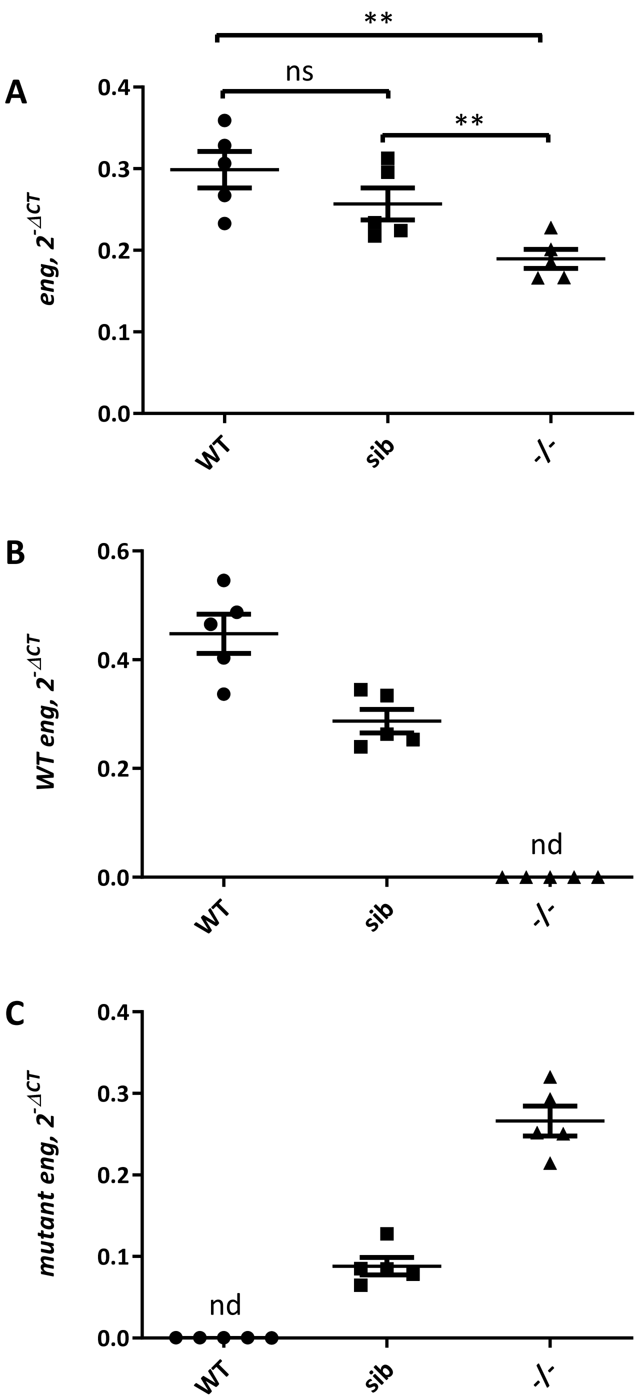
A
B
C
**
ns
**
eng, 2^-ΔCT
WT eng, 2^-ΔCT
mutant eng, 2^-ΔCT
0.0
0.1
0.2
0.3
0.4
0.6
WT
sib
-/-
nd

Fig. S1

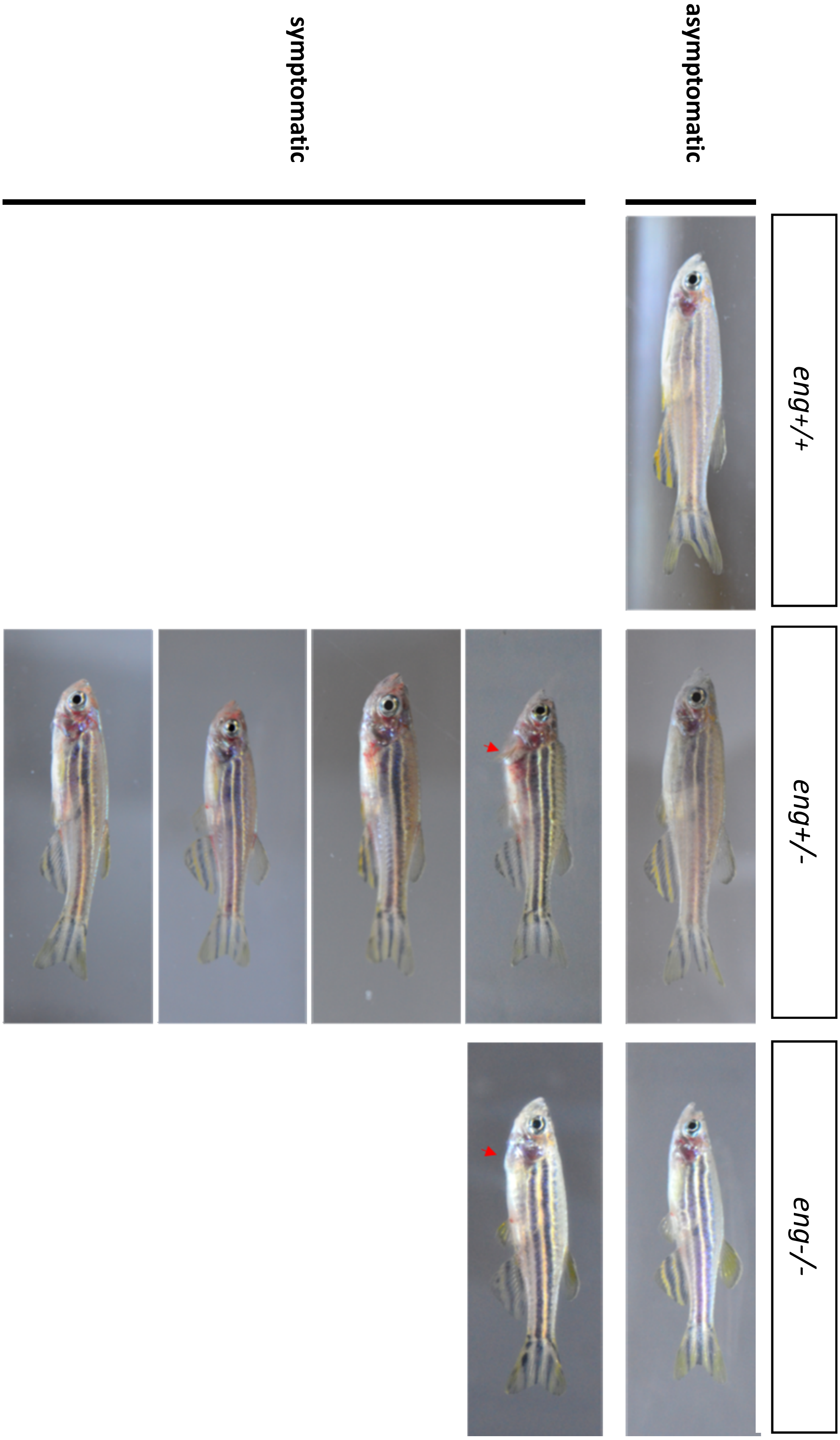


Fig. S2

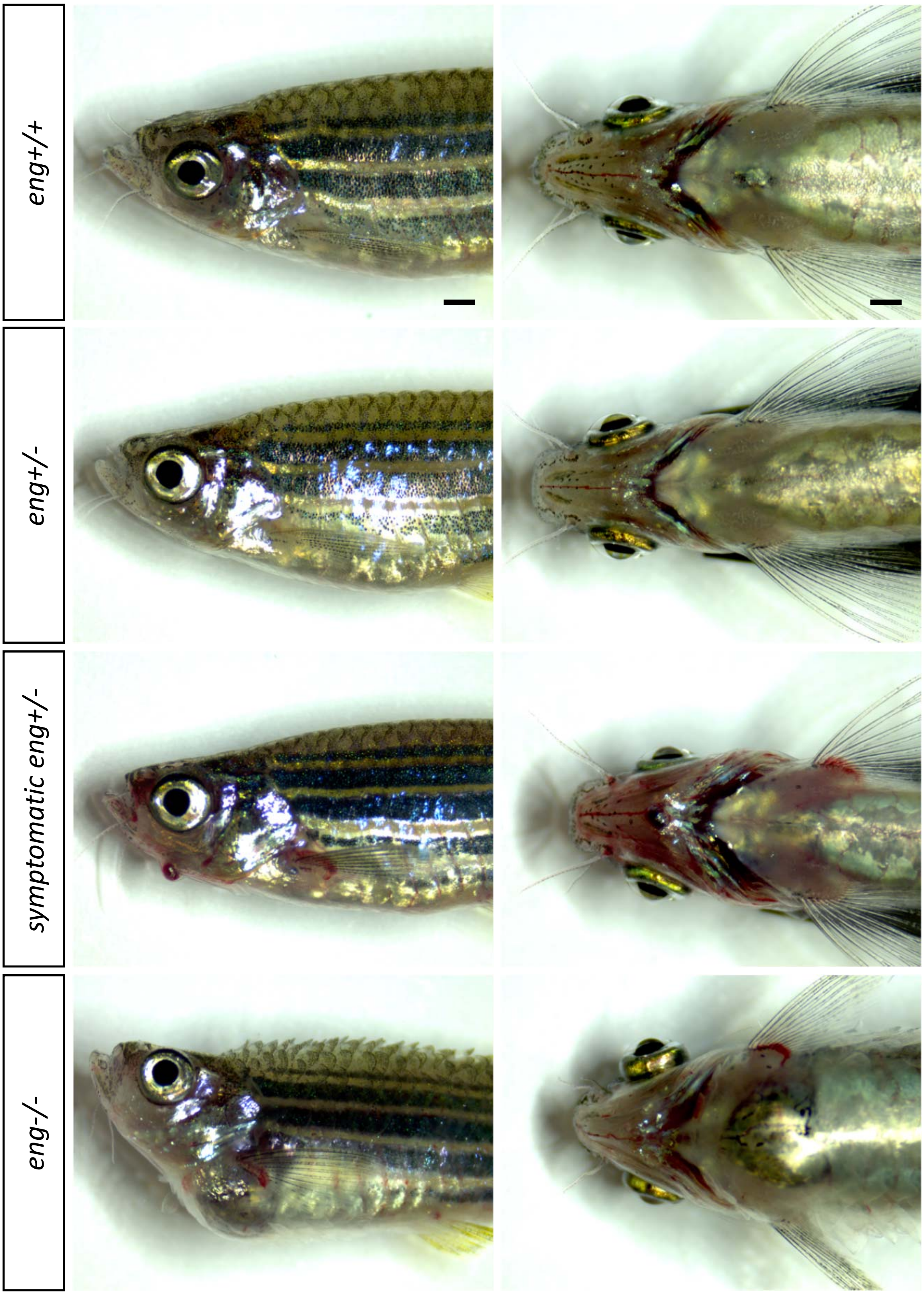


Fig. S3

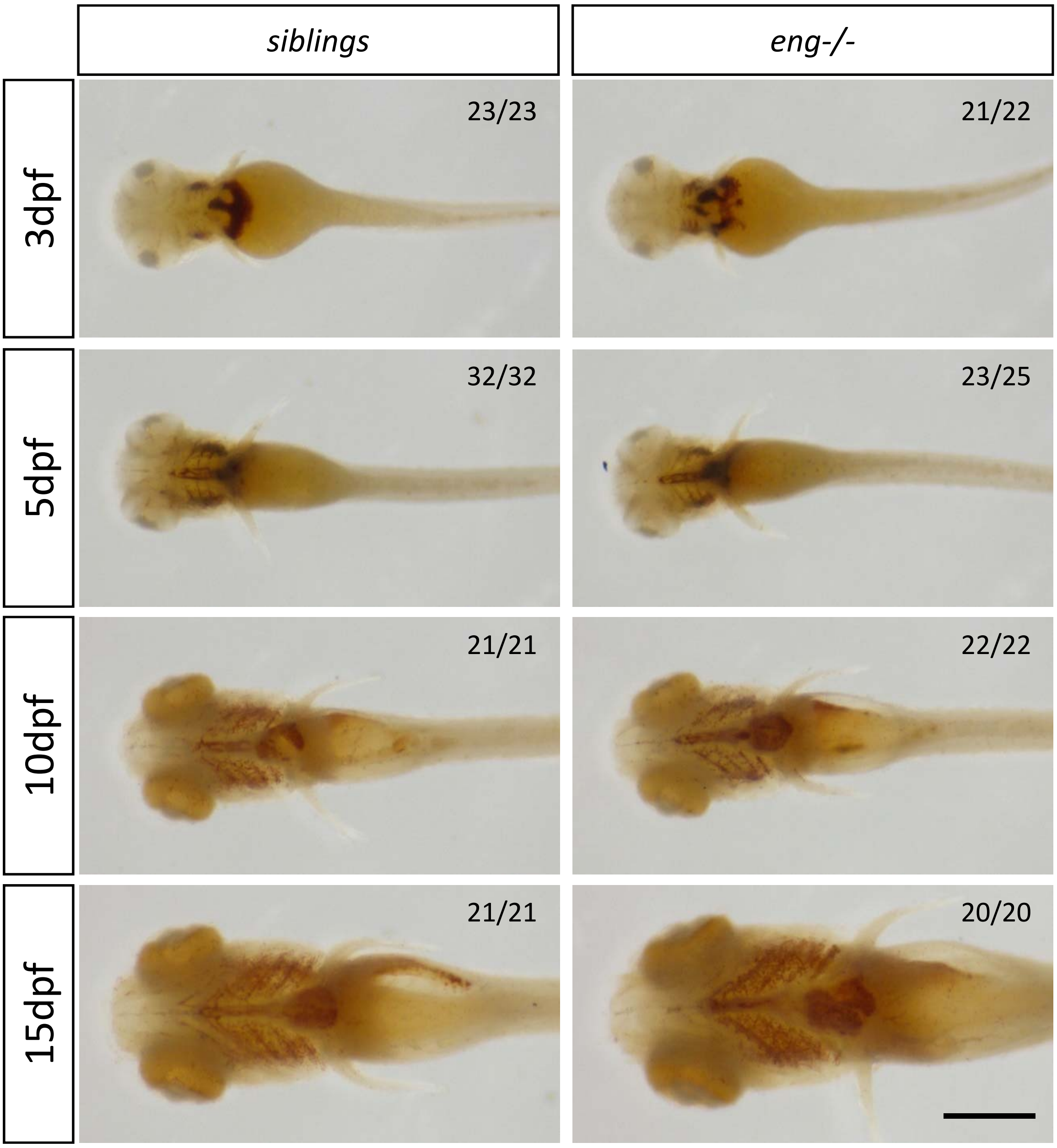
siblings
eng-/-
3dpf
23/23
21/22
5dpf
32/32
23/25
10dpf
21/21
22/22
15dpf
21/21
20/20

Fig. S4

**A**

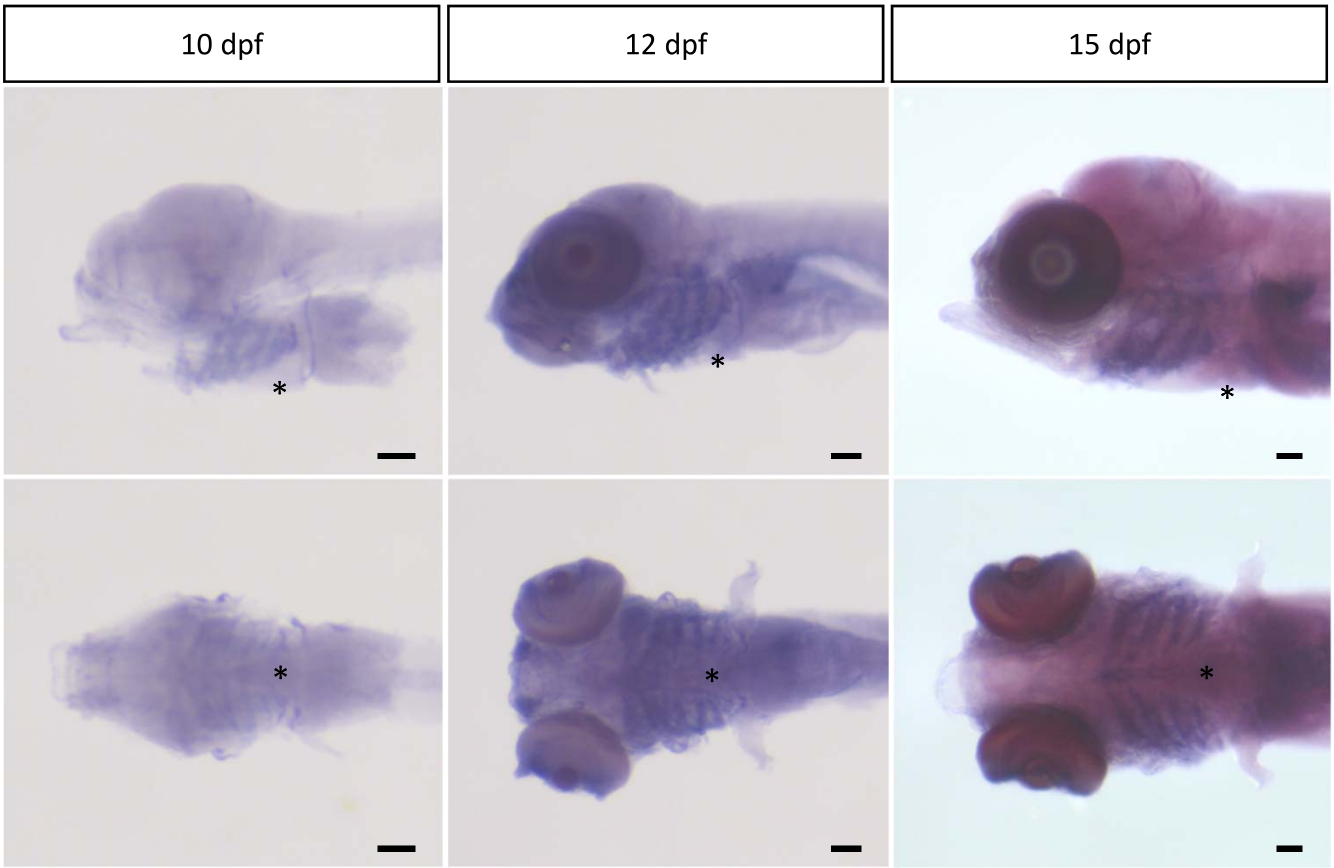


**B**

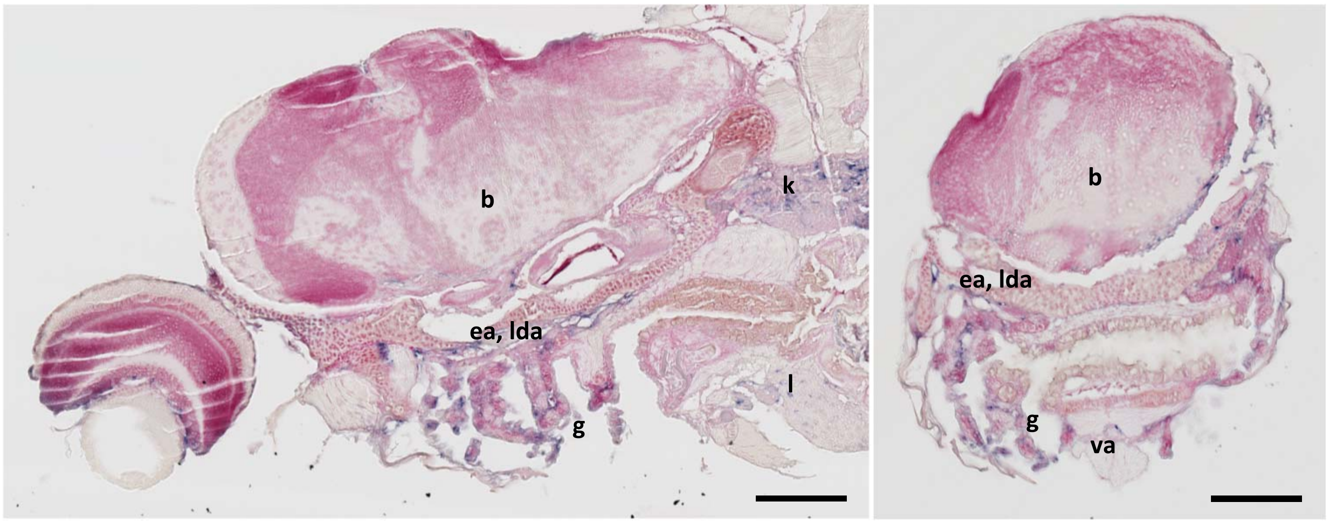


Fig. S5

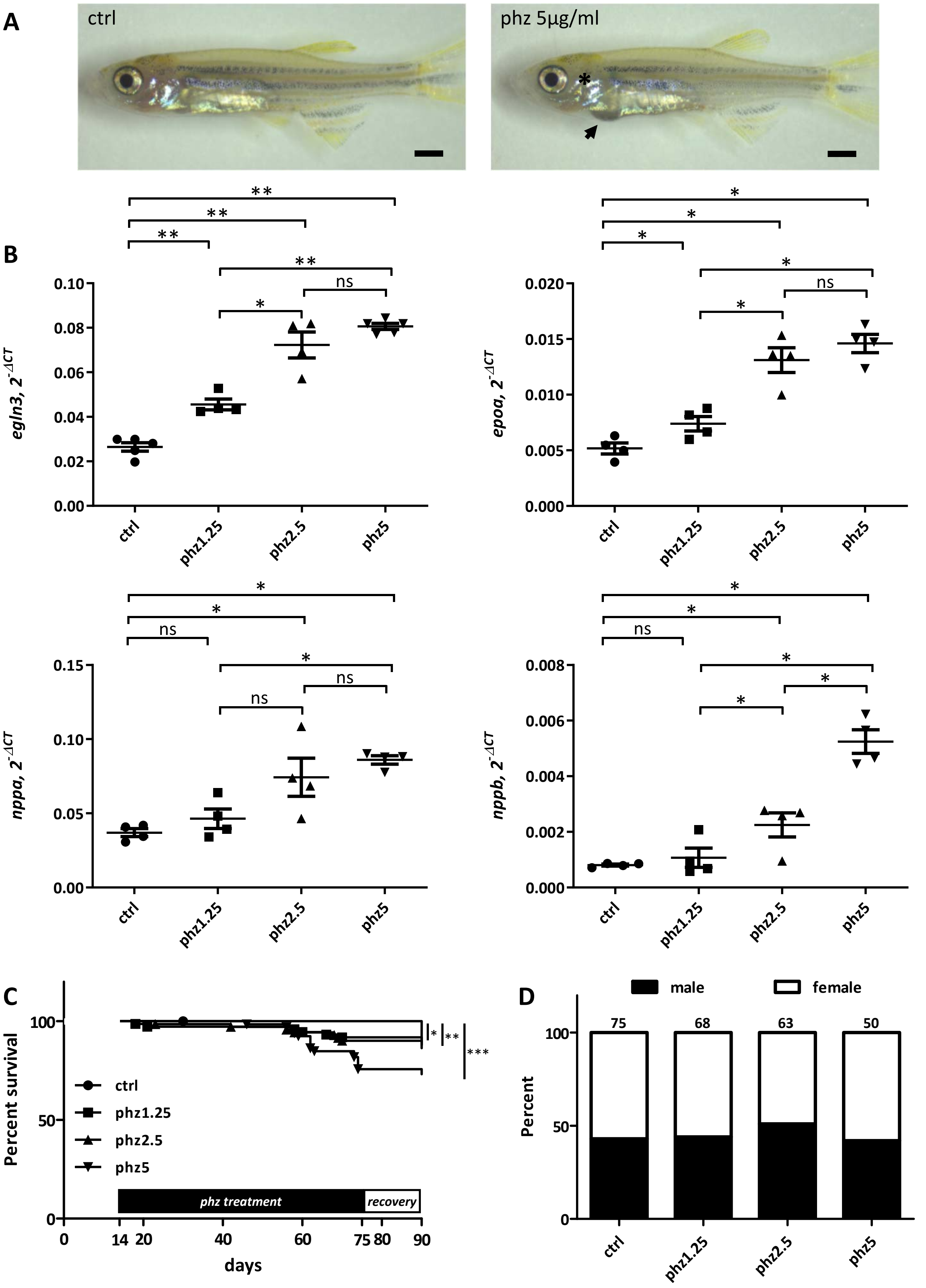
A
ctrl
phz 5µg/ml
*
B
egln3, 2^-ΔCT
epoa, 2^-ΔCT
nppa, 2^-ΔCT
nppb, 2^-ΔCT
ctrl
phz1.25
phz2.5
phz5
**
*
ns
C
Percent survival
ctrl
phz1.25
phz2.5
phz5
phz treatment
recovery
days
*
**
***
D
male
female
75
68
63
50
Percent

Fig. S6

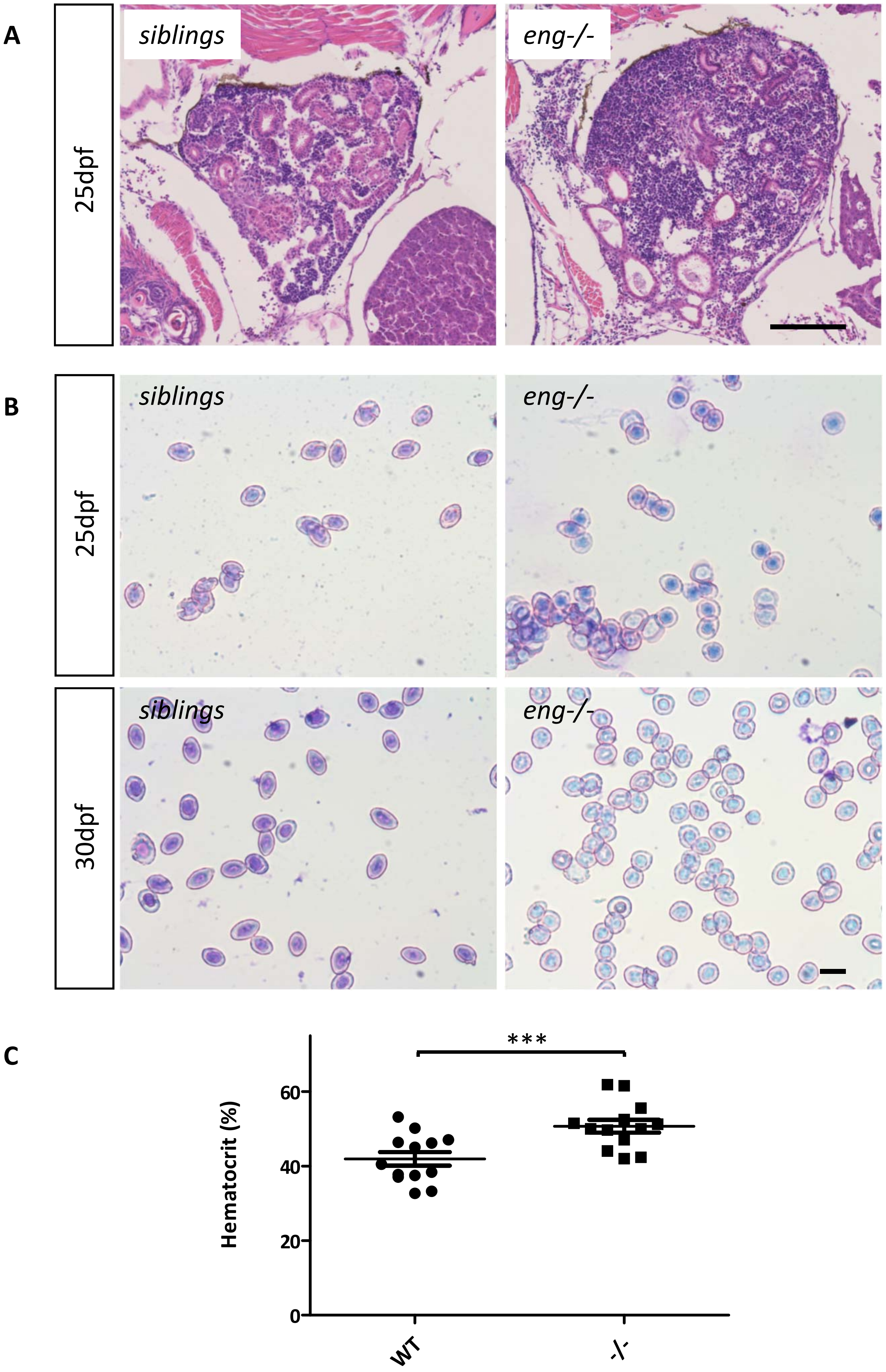

A
25dpf
siblings
eng-/-
B
25dpf
siblings
eng-/-
30dpf
siblings
eng-/-
C
***
Hematocrit (%)
60
40
20
0
WT
-/-


Fig. S7

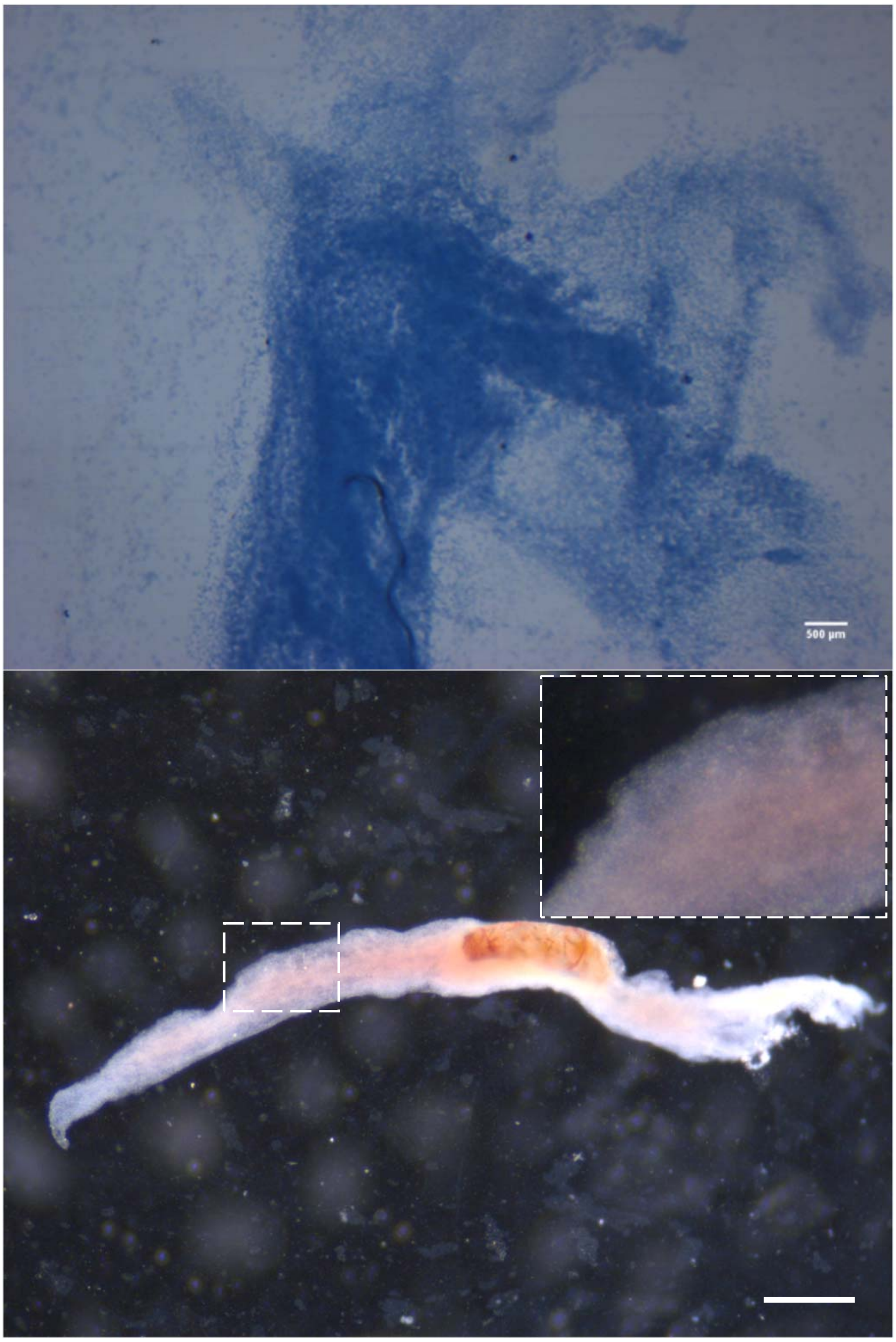


Fig. S8
